# Structure-affinity correlations and separable optical activity in carbon nanotube protein coronas

**DOI:** 10.64898/2026.07.16.739030

**Authors:** Jacob Miller, Alexis Coscia, Aceer Nadeem, Mijin Kim

## Abstract

Biomolecular corona on nanoparticle surfaces enriches low-abundance proteins and compresses the dynamic range of the plasma proteome. Single-walled carbon nanotubes are particularly attractive because their surface chemistry is highly tunable, and their near-infrared fluorescence directly reports corona modulation. However, how surface chemistry shapes the corona and how the corona relates to the fluorescence response remain unclear. We profiled protein enrichment patterns and the resulting fluorescence in a library of 25 chemically modified carbon nanotubes by quantitative proteomics and near-infrared fluorescence spectroscopy. We found that polymer and quantum-defect chemistry orthogonally modulate corona composition and dispersion forces are the dominant, universal driver of protein-nanotube binding. Proteomics and fluorescence correlation analyses indicate that enrichment and fluorescence response are partially decoupled. A subset of corona proteins stays optically silent, but where responses occur, enriched proteins respond at lower, physiologically relevant ranges than depleted ones. Our findings inform design principles for carbon nanomaterial-based sensors and biomarker discovery platforms.

---

The plasma proteome carries molecular signatures of cancer^1,2^, neurodegeneration^3^, and cardiovascular diseases^4^, making plasma proteins promising biomarkers for early diagnosis^5–8^. However, realizing this potential has been technically challenging. Protein concentrations in human plasma span more than ten orders of magnitude, with only the top 22 highest-abundance species accounting for roughly 99% of total protein mass.^9^ Additionally, the coexistence of thousands of distinct protein species complicates low-abundance protein analysis and biomarker discovery^10^. Conventional depletion and fractionation strategies^11^ partially address these issues^12^, at the cost of throughput and reproducibility^13^.

Nano-omics has emerged as a promising approach for enrichment and subsequent omics analysis of biofluids^14^. When introduced into plasma, a nanoparticle spontaneously absorbs a layer of proteins called a protein corona. A chemically diverse panel of nanoparticles can effectively compress the plasma’s dynamic range and fractionate proteins in parallel^15–17^. Among the possible nanoparticle candidates, single-walled carbon nanotubes (SWCNTs) have unique properties well-suited to nano-omics. Their surface chemistries are highly tunable through non-covalent polymer wrapping^18^ and covalent quantum well defects (QWDs)^19^, each of which imparts selectivity for specific analytes^20,21^. SWCNT fluorescence is sensitive to the local microenvironment^22–24^. A SWCNT-based nanosensor library combined with machine learning classified cancers^25,26^ from blood samples using differences in corona-driven fluorescence response. The SWCNT elements that significantly contribute to fluorescence-based target classification can be used in downstream corona analysis^25^, which increases throughput and precision for biomarker discovery. However, current SWCNT panel design is performed ad hoc, in which a random set of polymer and QWD types is combined to maximize surface orthogonality. Little work has explored the quantitative relationship between surface chemistry features^27,28^ and simultaneously connected corona composition and fluorescence transduction of SWCNTs.

Herein, we address both questions using a chemically diverse library of surface-engineered SWCNTs, characterized by quantitative proteomics and NIR fluorescence spectroscopy. Results suggest that polymer wrapping sets the dominant electrostatic and structural selectivity, while QWDs preferentially enrich compact, nonpolar proteins. Feature-level analysis of physicochemical protein descriptors reveals that dispersion forces drive protein-nanotube binding, outcompeting pi-pi contacts from surface-exposed aromatic residues, in contrast to previous understanding^29,30^. Pairing proteomics with fluorescence measurements shows a disparity between protein corona composition and resulting fluorescence response. Certain corona proteins remain optically silent regardless of enrichment, but where a response occurs, enriched proteins are more likely to respond at lower, physiologically relevant ranges rather than depleted proteins. This quantitative structure-affinity-activity relationship has important implications in rational surface engineering for next-generation sensors and biomarker discovery platforms.

## Results

### Protein corona extraction from a chemically diverse nanotube panel

We designed a nanotube panel as a product of 5 polymer types and 5 QWD types to represent diverse surface chemistry (**Fig 1a**). Non-covalent polymer wrapping included two PEGylated lipids (DSPE-PEG-Amine and DMG-PEG) and three DNA sequences ((AT)_15_, (GT)_15_, CT_2_C_3_T_2_C). The covalent QWD dimension comprised four QWNTs, including 4-carboxyl (carboxyl), 3,5-dinitro (dinitro), 3,4,5-trifluoro (trifluoro), and *N,N*-diethyl-4-amino (amino) aryl QWDs, with unfunctionalized (pristine) nanotubes as a reference. QWDs span a 5.7 log-unit hydrophobicity (logP) range and a charge at pH 7.4 from -0.999 to +0.126 (Supplementary Table 1).

**Figure 1.**
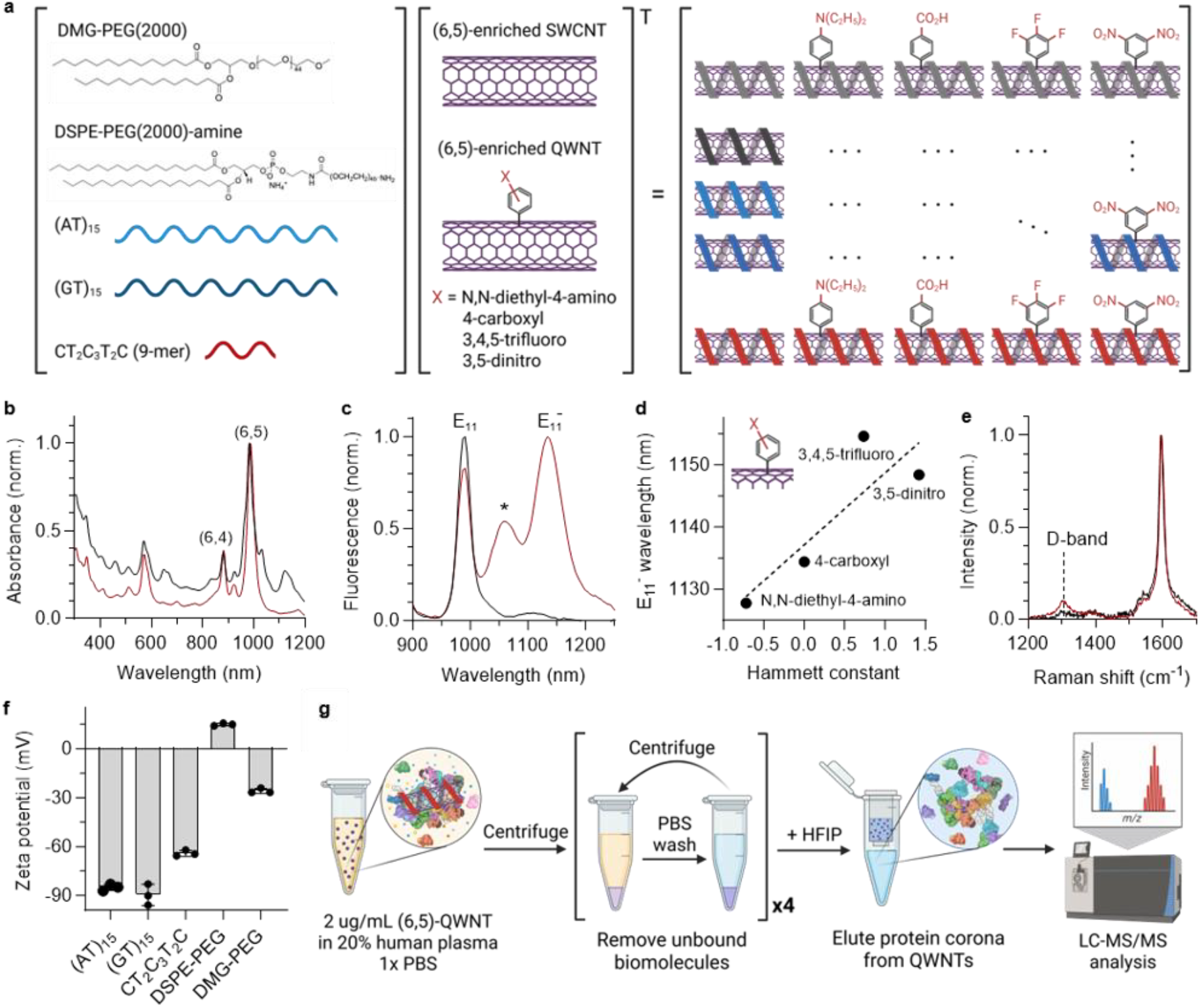
Synthesis and characterization of polymer-QWNT library for protein corona analysis. **a**, Schematic of library design. **b**, Absorption spectra of SG65i in 1% sodium deoxycholate (aq) before and after chirality sorting (black and red, respectively). **c**, Normalized fluorescence spectra of purified pristine (black) and N,N-diethyl-4-aminoaryl functionalized (6,5) nanotubes (red) in 1 wt% sodium deoxycholate (aq). The excitation wavelength is 575 nm. Asterisk (*) indicates (6,4) QWD emission. **d**, Correlation between the Hammett constant of the terminating group of QWD and the center wavelength of QWD emission. **e**, Normalized Raman spectra of pristine (black) and N, N-diethyl-4-aminoaryl functionalized (6,5) nanotubes (red). **f**, Zeta potential of (6,5) enriched nanotubes with N,N-diethyl-4-aminoaryl QWDs with different polymer wrappings. **g**, Schematic of protein corona extraction from plasma-incubated QWNTs with subsequent LC-MS/MS analysis.

Each polymer-QWNT construct was prepared by covalent functionalization of SWCNTs^31^, followed by (6,5) chirality sorting^32^. Aqueous two-phase extraction enriched (6,5) chirality up to 85%, evidenced by an increase in the relative intensity of the (6,5) absorption band (**Fig 1b**, Supplementary Fig. 1). Successful QWD functionalization introduced a new fluorescence peak (E_11_^-^) that is 100-200 meV redshifted from the intrinsic E_11_ emission (**Fig 1c**, Supplementary Fig. 2). The E_11_^-^ emission wavelength linearly correlated with the Hammett constant, a quantitative measure of the electron-donating/withdrawing capability of aryl QWD (**Fig 1d**)^33^. An increase in the Raman D-band to G-band ratio further verified the addition of sp^3^ defects on the nanotube sidewall and comparable defect density across QWD types^34^ (**Fig 1e**, Supplementary Fig 3). The resulting (6,5)-enriched QWNTs were resuspended in the polymer^35^, and their colloidal stability and surface charge were characterized with zeta potential (**Fig 1f**, Supplementary Fig. 4). We then extracted tightly bound corona proteins from the nanotubes after 2-hour incubation in pooled human plasma (**Fig 1g**, Supplementary Fig. 5).

### Polymer wrapping and covalent QWD chemistry independently shape the protein corona

LC-MS/MS identified a total of 1556 proteins, including 289 commonly identified proteins across all sensor coronas (Supplementary Fig 6). Principal component analysis (PCA) of the 289-dimensional protein fold-change (FC) matrix revealed that corona variance partitions along two orthogonal axes in the PC1-PC2 plane (**Fig 2a**). PEG- and DNA-wrapped nanotubes separate in one diagonal, and defect groups separate orthogonally. The most prominent QWD effect appears between pristine and QWNT coronas, while finer separation between defect types is still visible. The addition of PC3 (12%) separates the dinitroaryl QWD coronas from other polymer-QWNTs (Supplementary Fig. 7) presumably due to proteins with preferential affinities for non-(6,5) chirality^27^ in the dinitroaryl QWNTs (Supplementary Fig. 8). The combination of PC1-3 accounts for 60% of the total variance.

**Figure 2.**
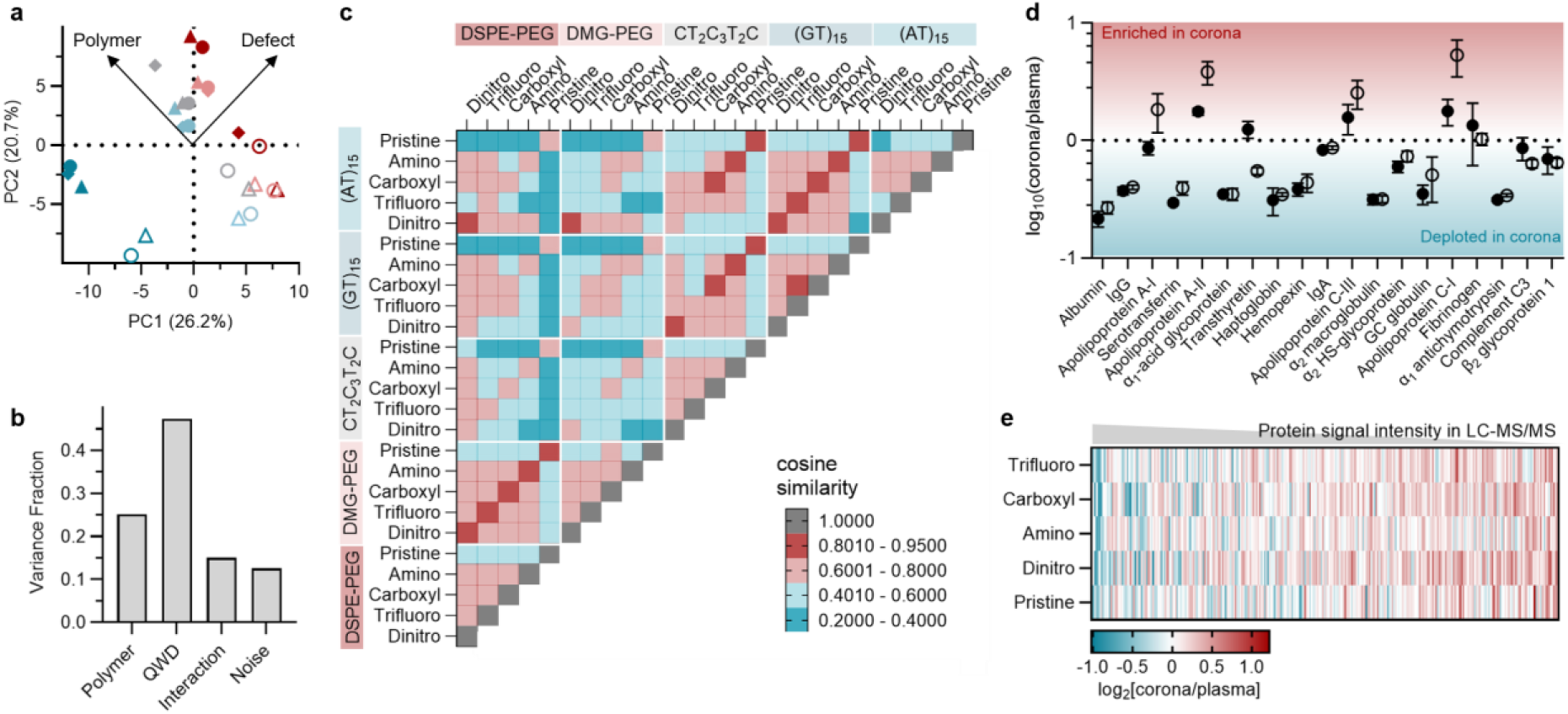
Effect of polymer wrapping and QWDs on protein corona composition. **a**, PCA plot of all sensor coronas, derived from Log2 protein fold-changes in 289 common proteins. Red, light red, gray, light blue, and blue symbols indicate dinitro, carboxyl, trifluoro, and aminoaryl QWNTs and pristine nanotubes, respectively. The filled triangle, circle, and diamond are CT_2_C_3_T_2_C, (GT)_15_, and (AT)_15_ wrapping, respectively. The empty triangle and circle are DMG-PEG and DSPE-PEG wrapping, respectively. **b**, Partitioned variance for library coronas for common proteins. **c**, Cosine similarity in corona compositions between QWNTs. **d**, Log10 fold-change in the top 19 most abundant plasma proteins by molarity. Average value across DNA (filled) or PEG (empty) wrappings. **e**, Heatmap of Log2 fold-change in protein abundance relative to plasma for (GT)_15_ wrapped QWNTs (ranked by plasma protein signal intensity). protein features, including global, structural, and surface-exposed features. Amino acid compositional features were reserved for independent validation.

To quantitatively assess the relative contributions of polymer and defect to corona composition, we fit a Multivariate Analysis of Variance (MANOVA) model to proteomic data with categorical predictors for polymer and QWD types, and their interaction (**Fig. 2b**). The model partitioned the total variance in the FC matrix into additive polymer (25%), additive QWD (47%), polymer-QWD interaction (15%), and residual components (13%). All terms were statistically significant. Refitting the model after excluding pristine nanotubes further balanced the effects of polymer and defect (Supplementary Fig. 9). The dominant additive effects over the interaction terms support the PCA observations that polymer and QWD are largely independent dimensions of corona differentiation.

The pairwise cosine similarity between their common protein fold-change vectors mirrors the PCA structure and MANOVA results (**Fig 2c**). In the polymer axis, DNA vs. PEG comparisons consistently yielded the most orthogonal pairings. QWD chemistry introduced a secondary, separable modulation from the polymer effect. The most notable QWD effect was between pristine and their QWNT counterparts.

We then interrogated the effect of corona adsorption on the plasma protein dynamic range. Among the 19 most abundant plasma proteins^36^, most were significantly depleted in the coronas relative to plasma, except for apolipoproteins (**Fig 2d**, Supplementary Fig. 10). For a global comparison, we arranged proteins by signal intensity in the plasma control tandem mass tag (TMT) channel (**Fig 2e**). While the signal intensity of a given protein is not perfectly proportional to its original sample concentration, comparisons can be made across broader groupings. The enrichment profiles indicate that the nanotube surface enriches lower-abundance proteins relative to the original plasma.

### Feature-level analysis of protein affinity to nanotube surface chemistry

We extracted 97 molecular features of all confidently identified proteins across the polymer-QWNT library and organized them into four categories: (1) global features describing the protein as a whole, (2) surface-exposed features reflecting solvent-accessible residues, (3) structural features representing fold and flexibility, and (4) compositional features capturing amino acid frequency profiles (Supplementary Table 2). Our initial analysis prioritized physically interpretable

First, within each QWD grouping, we analyzed Pearson correlations between individual protein features and their log_2_FC relative to plasma across all identified proteins (**Fig 3a**, Supplementary Fig 11). All coronas are positively correlated with flexibility, relative surface area, and relative enrichment in alpha helices, while negatively correlated with relative enrichment of beta sheet. These features suggest that dispersion forces play a significant role in protein binding to nanotube surfaces universally. Proteins with greater flexibility (more helices) and a larger relative surface area can maximize nanotube surface contact, increasing dispersion attraction force. Conversely, rigid proteins with beta sheets reduce their ability to adapt to the nanotube surface geometry. While dispersion forces are typically negligible compared to other intermolecular forces in small molecules, they scale with surface area and can dominate in protein binding.

**Figure 3.**
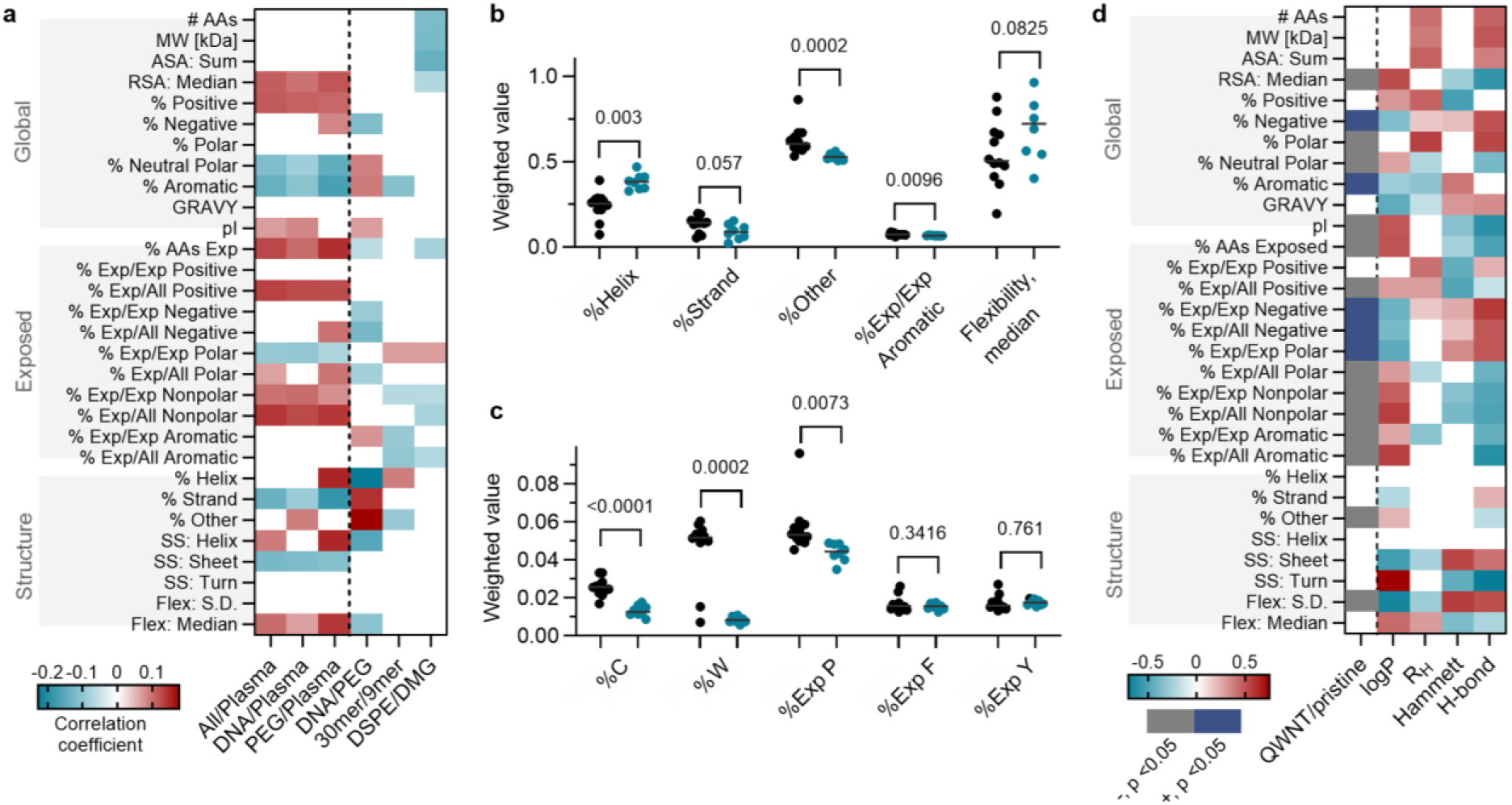
Quantitative structure affinity relationships of the protein corona. **a**, Pearson correlation heatmap linking weighted protein corona features to polymer wrapping properties with respect to plasma (left) and within nanotube coronas (right). Only p-values of <0.05 are shown. **b**, t-tests of flexibility related macro features of DNA (black) and PEG (blue) wrapping. **c**, t-tests of flexibility related amino acid level features of DNA (black) and PEG (blue) wrapping. **d**, Effects of QWDs on protein corona enrichment. First left column: t-test heatmap linking weighted protein corona features to polymer wrapping properties. The gray-and-blue color bar indicates significant features with negative and positive correlations, respectively. Only p-values of <0.05 in unpaired, nonparametric Mann-Whitney tests are shown. Remaining columns: Spearman correlation heatmaps linking weighted protein corona features and QWD chemical parameters. Only p-values of <0.10 are shown.

Interestingly, nearly all nanotubes showed a significant negative correlation with % Aromatic. This contrasts with the previous understanding, which suggested that pi-pi binding between aromatic amino acid residues and the nanotube surface should strongly drive adsorption^29,30^. These prior reports are derived from free amino acids or short peptides, in which all residues are solvent accessible. However, in proteins, aromatic amino acids often pi-pi stack with each other in the hydrophobic core^37,38^, preventing their interaction with the nanotube surface, as well as increasing protein rigidity^38,39^. The reduction in dispersion forces from pi-pi stacking within proteins likely lowers protein affinity. This is further confirmed by the fact that % Exposed Aromatic does not show statistical significance relative to plasma.

### Polymer-dependent structural selectivity of protein corona

We calculated log_2_FC values for the average of polymer groups relative to each other (last three columns of **Fig 3a**). Between PEG and DNA, electrostatic and dispersion effects were significant. The surface charge of the nanotubes differentiates protein affinities, with the isoelectric point (pI) of proteins positively correlated with DNA/PEG. As the pI of a protein increases, it will be more positively charged at neutral pH. Therefore, the densely negatively charged DNA-SWCNTs will recruit positively charged amino acid residues. This electrostatic term is further confirmed by the observation that PEG/Plasma positively correlates with both negatively and positively charged residues (hydrophilic interactions), whereas DNA/Plasma only positively correlates with positively charged residues. PEG and DNA are also differentiated by their affinity to flexible proteins. While both polymer types enrich flexible proteins relative to plasma, this enrichment is more pronounced in PEG-SWCNTs. This may reflect a greater loss of conformational entropy in the PEG layer upon protein insertion than in the rigid DNA layer on the nanotubes.

To extend our analysis to cross-defect comparison, each corona was collapsed into a single descriptor per feature (see Methods). At the macro level, prominent differentiations between DNA and PEG coronas lie in secondary structure and conformational flexibility (**Fig. 3b**). PEG wrapping enriches helical proteins with higher baseline flexibility. DNA wrapping enriches relatively rigid proteins with higher strand contents, disordered structure (% Other), and surface-exposed aromatic residues. The macro-level trends are consistent with the Pearson correlations of Fig 3a.

The global trends are further supported by amino acid-level weighted features (**Fig 3c**). DNA coronas are significantly enriched in cysteine (%C), consistent with recruitment of disulfide-stabilized, structurally rigid proteins. Tryptophan is also enriched in DNA coronas (%W) with a large aromatic side chain due to pi-pi stacking at the exposed nanotube sidewall. Exposed proline (% Exp P) is a rigid, helix-breaking residue and is enriched in the DNA coronas. No significant differences were observed in exposed phenylalanine and tyrosine.

### Effect of QWDs on protein corona composition

The most significant difference for QWD effects is between QWNTs and pristine controls (First column in **Fig 3d**). QWD introduction shifts the corona toward proteins with polar, negatively charged surfaces and compact, rigid folds. Exposed negatively charged residues are significantly enriched, and the median weighted pI drops from 8.26 to 7.23. Exposed nonpolar residues are depleted while exposed polar residues are enriched. These surface chemistry preferences are coupled with a consistent structural signature of compact, rigid folds. Surface exposure is reduced across multiple metrics (%AA Exposed, RSA Median), disordered content is depleted (% Other), and backbone flexibility variance is lower. Similarly, while total aromatic content is enriched, exposed aromatic residues are depleted, indicating that the aromatics reside in buried hydrophobic cores. Residue-level features independently supported polarity, charge, and rigidity selection (Supplementary Fig 12).

Spearman correlation between log_2_FC-weighted protein features and QWD descriptors resolved how the physicochemical properties of QWDs relate to corona composition (**Fig. 3d**, Supplementary Fig 13; descriptor-resolved correlations in **Supplementary Note 1**). Five chemical descriptors of the aryl QWD were considered with the following physicochemical categories: electronic (Hammett constant), steric (hydrodynamic size), and thermodynamic (H-bond capacity, logP). To distinguish genuine descriptor-specific effects from collinearity artifacts, we focus below on features exclusively significant for one descriptor class.

The electronic character of QWDs correlates with the depletion of positively charged surface residues. More H-donating and hydrophobic QWDs negatively correlate with GRAVY and positively with exposed aromatics, indicating a preference for hydrophilic proteins with high surface exposed aromatics. Structural features, including flexibility and secondary structure information, are significant for both Hammett and the thermodynamic axis, with opposite correlations. However, the two are collinear in the current library and cannot be deconvolved. In terms of steric space, bulkier QWDs recruit larger, more polar corona proteins, but do not significantly correlate with any structural or flexibility metric. This differentiates the steric axis related to protein size and polarity selection from the conformational effects captured by the electronic/thermodynamic gradients.

### Distinct drivers of variance in corona composition and fluorescence response

To examine whether the compositional differences in coronas translate into fluorescence modulation, fluorescence spectra changes upon plasma incubation were quantified (**Fig. 4a**). The E_11_ and E_11_^-^ spectral changes show clear grouping by polymer type (**Figs 4b-c**). The magnitude of the intensity changes scales with polymer packing density on the nanotube surface. CT_2_C_3_T_2_C (9-mer, most exposed sidewall) shows the greatest increase in intensity, followed by the 30-mer sequences (AT)_15_ and (GT)_15_. PEG wrapping showed minimal change to a decrease in intensity. In permutation-based variance partitioning of fluorescence response, each spectral feature is dominated by the additive polymer term, while the additive QWD effect wnas not statistically significant for any of the spectral features (**Fig 4d**).

**Figure 4.**
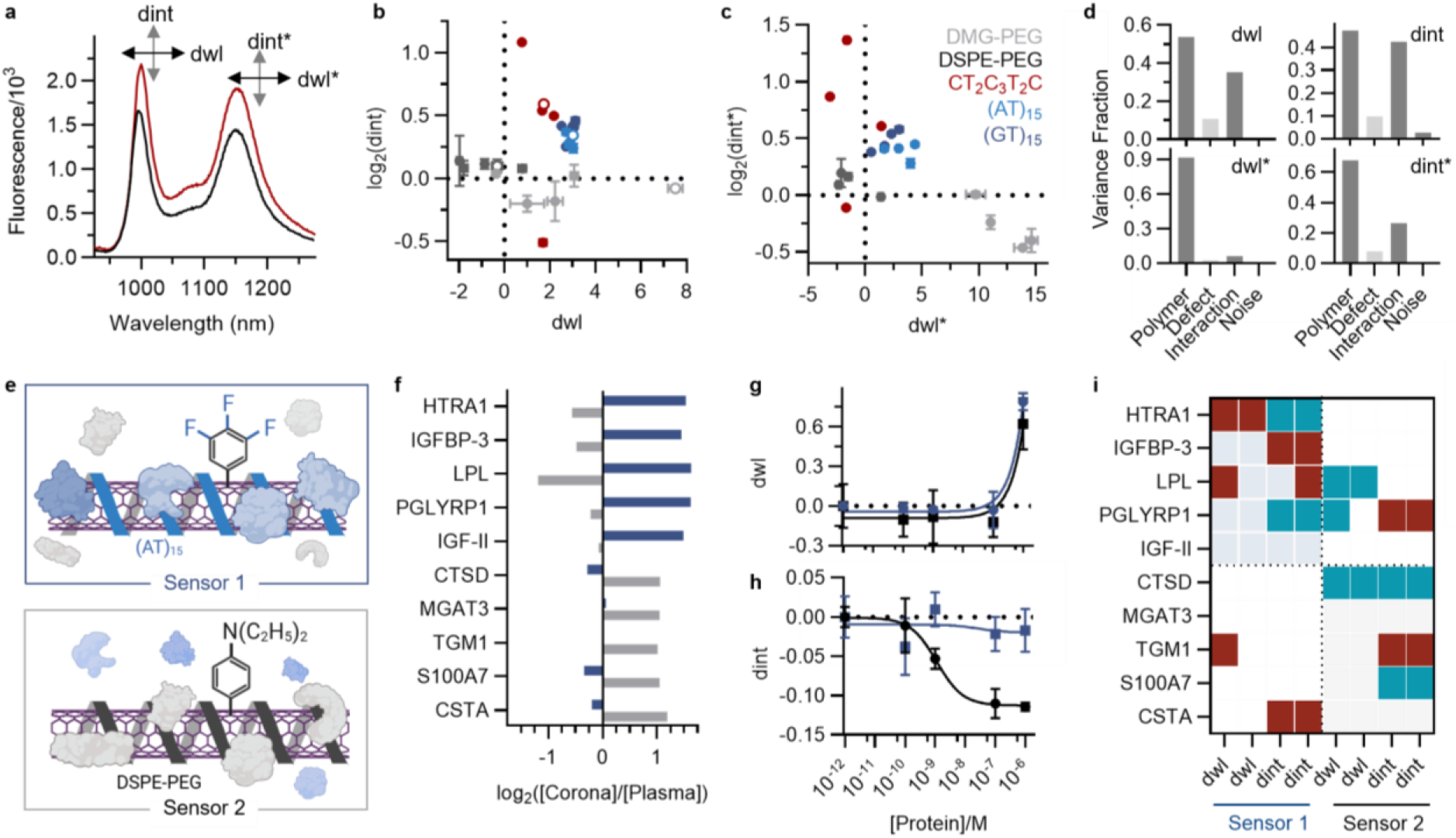
Quantitative structure-affinity-activity relationships of the protein corona. **a**, Representative fluorescence spectra of (GT)_15_ wrapped 4-carboxyaryl QWNTs in PBS (grey) and after two-hour incubation in 20% plasma (red). **b**, Correlation between intensity and wavelength changes of **b**, E_11_ and **c**, E_11_^-^ fluorescence bands across the complete polymer-QWNT array. Error bars are the standard deviation of triplicates. Open circles of **b** are unfunctionalized carbon nanotubes. Different colors denote polymer wrapping types. **d**, Partitioned variance for spectral parameters. Light gray indicates not statistically significant. **e**, Schematic of preferential protein enrichment on sensors 1 (top) and 2 (bottom) surfaces. **f**, Log2 enrichment with respect to plasma for all ten validation proteins on sensors 1 (blue) and 2 (grey). Representative concentration-dependent fluorescence responses for **g**, HTRA1 and **h**, S100A7. **i**, Heatmap of significant sensor responses across ten proteins and four spectral parameters per sensor. Color indicates response direction (blue: negative, red: positive).

The disparity between variance driving corona composition and variance driving fluorescence response suggests that a subset of corona proteins is unreported by fluorescence. To confirm the existence of this fluorescently silent protein corona, we designed an in-plasma titration experiment using purified proteins. Based on the cosine similarity analysis of Fig 2c, (AT)_15_ wrapped aminoaryl QWNT (sensor 1) and DSPE-PEG wrapped trifluoroaryl QWNT (sensor 2) were selected as a maximally orthogonal polymer-QWD pair (**Fig 4e**). From the proteomics data, we identified 10 proteins that were exclusively enriched for each sensor surface (**Fig 4f**, Supplementary Table 5). Each protein was titrated into 20% plasma from 1 pM to 1 µM, and the fluorescence features were tracked (Supplementary Fig 14). A response was classified as active if the spectral change significantly deviated from the baseline only at the highest concentration (Welch’s t-test; **Fig 4g**) or monotonic across concentrations as quantified by Spearman correlation (**Fig 4h**). The remaining features are classified as silent (inactive).

Within twenty protein-sensor pairs, enriched pairs were active slightly more often than the non-enriched pairs (7/10 vs. 4/10, **Fig. 4i**). A subset of strongly enriched proteins did not elicit significant spectral responses on either sensor (IGF-II, MGAT3), while several depleted proteins responded, violating a simple affinity-to-activity correspondence. However, where enrichment and response coincided, the responses that emerged at physiologically relevant, lower concentrations (pM-nM) were generally with enriched proteins, whereas responses to the non-enriched proteins often appeared only at the highest concentrations or not at all (Supplementary Fig 15). Lastly, we cross-referenced the corona enrichment of trifluoroaryl QWNT with DSPE-PEG wrapping (this work) against an independent fluorescence dataset (Goerzen et al^25^) and validated the observed enrichment-response relationship (Supplementary Figs 16-18).

## Discussion

We characterized protein corona formation across a combinatorial library of 25 polymer-QWNTs and established physicochemical rules governing differential protein enrichment from human plasma. Polymer wrapping and covalent QWD chemistry independently shape corona composition through broadly orthogonal mechanisms, providing a rational basis for sensor panel design. Importantly, corona enrichment shapes whether and how readily a protein modulates fluorescence but does not predict it one-to-one, and composition and optical transduction are only partially coupled.

At the feature level, dispersion forces are the universal, surface-independent driver of protein-nanotube binding. Proteins with greater conformational flexibility and accessible surface area are preferentially enriched in the coronas. This result is mechanistically coherent because larger, more flexible proteins maximize nanotube contact area and amplify cumulative van der Waals interactions at the macromolecular scale in ways that rigid, compact proteins cannot^40^. Polymer identity adds an electrostatic selection layer, in which negatively charged DNA wrapping preferentially recruits basic proteins through charge complementarity. Notably, the near-universal depletion of total aromatic amino acid content in corona proteins directly challenges the widely cited pi-pi stacking model of protein-CNT affinity derived from free amino acids or short peptides^29,30^. In intact plasma proteins, aromatic residues are predominantly buried in hydrophobic cores and structurally rigidify the fold, opposing the flexibility-dependent binding mechanism that dominates at the full-protein scale.

More importantly, we confirmed separable corona composition and fluorescence responses of carbon nanotubes. The fluorescence responses of polymer-QWNTs indicate the existence of fluorescently silent corona proteins and suggest a spatial separation between the protein’s surface binding site and the fluorescence-modulating binding sites on the nanotube surface. In addition, enriched proteins resolved at lower concentration more often than depleted proteins. This partial link between affinity and activity suggests that the molecular features setting corona affinity also shape transduction, and that a quantitative structure-affinity-activity relationship could be learned to predict sensor response from physicochemical properties of proteins.

The independence of polymer and QWD effects on corona composition has direct implications for sensor panel design. Polymer identity establishes the dominant corona identity, with DNA- and PEG-wrapped nanotubes producing the most orthogonal corona profiles across the library. QWD chemistry introduces a secondary, separable modulation within each polymer class. This is consistent with earlier reports on machine perception-based disease classification, in which either QWD or polymer type was the unifying determinant of diagnostic performance^25,26,42^. In practice, this argues for assembling the initial array by selecting nanotubes whose corona-enrichment profiles are maximally orthogonal so that the panel collectively samples the broadest slice of the proteome and response coverage. Because corona composition does not map onto fluorescence with one-to-one correspondence, each candidate sensor should be benchmarked for response diversity such that the array is diverse in both composition and transduction. Finally, it is possible that fluorescence-based screens systematically miss a subset of corona enrichment, necessitating panel design to optimize for maximal diversity in fluorescence sensitivity.

Future work that systematically compares features across the silent and active protein classes will be necessary to develop accurate predictive models of optical transduction competence for identifying biomarker candidates. In addition, expanding the corona characterization to metabolomic dimension^43^ would capture in-depth compositional complexity of the biomolecular corona and illuminate how proteins, lipids, and small molecules interact, which is fundamental to understanding the physicochemical rules that determine corona composition in nanomaterials^44^.

## Software and Code Availability

All statistical analyses were performed in Python 3.12 using pandas (v2.0), NumPy (v1.24), and SciPy (v1.11). Spectral data processing was conducted with custom MATLAB codes (R2024b). Figures were generated using GraphPad Prism (v11.0.0) and Biorender. Custom analysis scripts are available at [repository URL/To be shared].

## Author contributions

**M.K**. conceived the article. **J.M**. and **A.C**. conducted the experiments. **J.M**. and **M.K**. analyzed data for the article. **J.M**., **M.K**., and **A.N**. wrote, reviewed, and edited the manuscript before submission. All authors have given approval to the final version of the manuscript.

## Acknowledgments

This work was supported in part by the National Institutes of Health (R00-EB033580) to M.K. This research was funded, in part, by the Advanced Research Projects Agency for Health (ARPA-H) under Agreement No. 1AY2AX000080-01. The views and conclusions contained in this document are those of the authors and should not be interpreted as representing the official policies, either expressed or implied, of the U.S. Government. J.M. was supported in part by the NSF GAANN Fellowship. The proteomics data were acquired at the Systems Mass Spectrometry Core Facility at the Georgia Institute of Technology, funded by the NIH (1S10OD038327-01). Raman data were acquired at the Materials Characterization Facility of the Institute for Matter and Systems at the Georgia Institute of Technology. We thank Dr. Dana Goerzen for sharing raw fluorescence data from his published work and for helpful feedback.

## Methods

### Solid state synthesis of quantum well defect modified carbon nanotubes

Raw SG65i (Sigma-Aldrich, Cas-No: 308068-56-6) were covalently functionalized with previously reported solid-state functionalization protocol by Piletsky et al^31^ Briefly, raw SWCNT soot was combined with phosphate buffered saline (PBS, Sigma-Aldrich, Lot: 0000291288) and diazonium salts or anilines (detailed reactant ratios are provided in Supplementary Tables 3 and 4) and briefly vortex mixed. After incubation at room temperature for specified time, the reaction mixture was transferred to a 5 mL syringe (Henke-Ject) with a 20 µm polyethylene frit (Sigma-Aldrich). The reaction solution was ejected, and the remaining SWCNT powder was thoroughly rinsed with water (2 × 5 mL), 70% ethanol (2 × 5 mL), and water again (2 × 5 mL). CNT powder was then added to aqueous solution of sodium deoxycholate (1 w/w%, >97%, Sigma Aldrich) to adjust the final concentration of 0.8 mg/mL. Nanotubes were suspended via tip sonication (Fisherbrand Model 120 Sonic Dismembrator) at 70% amplitude and 4 °C for 30 min. The sonicated nanotube solutions were centrifuged at 21,300*g* for 30 min at 20 °C. The top 80% of the supernatant was collected for further chirality sorting.

### Chirality enrichment of QWNTs by aqueous two-phase extraction

(6,5) chirality was enriched using a surfactant-based aqueous two-phase extraction^32^. Stock 1 is prepared by combining 42.3 g of 6 kDa polyethylene glycol (Thermo Scientific, Cat: A17541.0B, Lot. 10253547), 14.7 g of 70 kDa dextran (TCI, Cat: 9005-54-0, Lot. 7ALSL-MS), 1.47 g of sodium cholate (Sigma-Aldrich, Lot: 102640683), 3.351 g of sodium dodecyl sulfate (Sigma-Aldrich, Lot: 0000286470, ≥98.5%, 0.426 g of sodium chloride (Sigma-Aldrich, Lot: 0000275233, ≥99.0%), and 300 g of nanopure water. Stock 1 is mixed overnight at room temperature to ensure proper dissolution. The following day, 800 µL of CNT dispersion was combined with 1.581 mL of water and 15.18 mL of Stock 1. Mixture was thoroughly vortexed and centrifuged at 6000*g* for 5 min in a fixed-angle centrifuge (Eppendorf 5430R). The resulting top phase and interface were aspirated off, and the bottom phase was diluted 1:20 in 1 wt% sodium deoxycholate (aq). Purification was performed using dialysis membrane centrifugal filters (Amicon Ultra, Regenerated Cellulose, 100K NMWL, 15 mL, Lot: 0000433432). 15 mL of the diluted bottom phase was loaded onto the centrifugal filter and filtered at 2000*g* for 5 min. The process was repeated until a volume of 1 to 1.5 mL remained, thoroughly resuspending filter-bound nanotubes after each 5 min pulse. This filtration process was repeated for at least 3 cycles to reduce the polymer concentration by a factor of 10^3^. After 3 cycles, a small amount of CNT suspension was removed and combined with methanol (≥99.9%, Sigma-Aldrich, Lot: 1003796521) in a 1:2 volumetric ratio. The mixture was briefly vortexed and examined for turbidity resulting from residual polymer contamination. If turbidity was present, the purification cycle was repeated, and the sample was tested with methanol again. Once the CNT/methanol mixture was no longer turbid, the CNT suspension was examined using UV-vis-NIR absorption spectroscopy to quantify (6,5) concentration. After purification and concentration, functionalized CNT stock solutions were stored in the dark at room temperature.

### Polymer Rewrapping

Single-stranded DNA from Integrated DNA Technologies, DSPE-PEG 2000 (Avanti, Lot: 880128P-50MG-A-077), DMG-PEG 2000 (Avanti, Lot: 880151P-5G-A-027) were used as received. Polymer stock solutions were prepared as follows. For ssDNA, DNA was combined with DI water and pipetted thoroughly to form 10 mg/mL solutions. For modified PEG, the dry polymer powder was combined with water and bath-sonicated for 30 min (37 kHz, 60% power) to form a 5 mg/mL solution. To exchange the surfactant wrapping on the nanotube surface with the desired polymer wrapping agent, the following protocol^35^ was used. In a 2 mL Eppendorf tube, 250 µL of the selected CNT stock was combined with 50 µL of either ssDNA or PEG stock solution. While vortexing at low speed, 1 mL of methanol was added slowly to precipitate nanotubes, followed by 50 uL of 5 M NaCl. Precipitated nanotubes were then removed from vortexing and centrifuged at 21,300*g* for 5 min. After centrifugation, the supernatant was aspirated and 250 µL of DI water was added. The CNT precipitate was then vigorously bath sonicated and pipetted until pellet was fully resuspended. The cycle of polymer addition, precipitation, and resuspension in water was then repeated once more. After the second rewrapping, the polymer-wrapped nanotube suspension was then combined with 100 µL of additional polymer stock and diluted with 650 µL of DI water to form a 1 mL suspension of 1.0 mg/mL DNA or 0.5 mg/mL PEG, respectively. The resulting 1 mL suspensions were then tip sonicated at 30% power for 30 min.

After tip sonication, polymer-wrapped CNT suspensions were loaded into a floating dialysis membrane (Float-A-Lyzer G2, MWCO: 1000 kD, 1 mL) and placed in 1 L of nanopure water. Dialysis was performed overnight with bath exchange at 1 hr and 4 hr at room temperature. After dialysis, suspensions were characterized by UV-vis-NIR absorption, fluorescence, and zeta potential. Dialyzed nanotube suspensions were used in plasma incubation the same day as dialysis completion.

### Plasma Incubation and Protein Corona Extraction

Pooled human plasma (Blood derived, K2 EDTA, Innovative Research, Inc.) was thawed and centrifuged at 13,000*g* at 4°C for 10 min. After centrifugation, the resulting fatty surface layer was aspirated, and 600 uL of plasma supernatant was combined with 300 uL of 10x PBS, polymer-wrapped CNT stock, and water to form a 3 mL mixture of (6,5) CNT solution with the optical density at (6,5) E_11_ of 0.15. The resulting (6,5) nanotube concentration is approximately 2 µg/mL. The resulting mixture was placed on a Fisherbrand Multi-Platform Shaker in a closed container at 200 speed for 120 min. The incubated mixture was divided into 3 × 1 mL aliquots in Eppendorf Protein LoBind Tubes (1.5 mL, Lot: O221182K) and centrifuged at 21,300*g* at 4°C for 25 min. After centrifugation, a visible CNT pellet appears at the bottom of the container. The supernatant is gently aspirated off and pellets are recombined by transferring 1250 µL 1X PBS between the 3 aliquot tubes to collect the pellets into a single Eppendorf tube. This process is repeated two more times. For final resuspension, pellets are suspended in 200 uL of 1,1,1,3,3,3-hexafluoro-2-propanol (HFIP) (>99%, Sigma-Aldrich, Lot: 1003743061), transferred to a PTFE membrane filter tube (Ultrafree-MC Centrifugal Filters, MilliporeSigma, 200 µm, Cat: UFC30LG25, Lot: N5BB27060), and centrifuged at 12,000*g* for 10 min. The HFIP elution step is repeated once more, resuspending the nanotube pellet in the supernatant chamber and centrifuging. Protein-containing HFIP elution is transferred to a fresh Protein LoBind tube, using additional HFIP to thoroughly remove all proteins from the filter tube. The protein suspension was then stored at -80°C until digestion.

### Calculation of Weighted Protein Features

Amino acid-derived information and physicochemical descriptors, e.g., hydrophobicity index (GRAVY), were computed from primary protein sequences using BioPython^45^. Residue-level structural annotations, including relative solvent accessibility, secondary structure propensity, etc., were predicted using NetSurfP^46^. To enable direct comparison across all 25 constructs, we collapsed each corona into a single descriptor per feature by summing the product of each protein’s fold change relative to plasma and its corresponding feature value across all identified proteins and dividing by total weight:

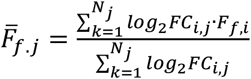

where N_j_ is the total number of proteins identified in that corona; FC_i,j_ is the abundance ratio of protein i in the nanotube construct, j; F_f,i_ is the value of feature f for protein i. This yields a single enrichment-weighted descriptor per nanotube construct, 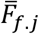, representing the average physicochemical character of the corona, where highly enriched/depleted corona proteins contribute proportionally larger positive or negative weighting, respectively.

### Spectroscopic Characterization

For the absorption and fluorescence measurements, the nanotubes were dispersed in an aqueous solution of polymers or sodium deoxycholate as described in the earlier paragraphs. All measurements were taken in Eppendorf UVette Cuvettes (Cat: 952010051). Absorbance spectra of nanotubes were collected by a V-780 UV-Visible-NIR spectrophotometer (Jasco, Inc.) with the following parameters: UV-Vis bandwidth of 2.0 nm, NIR bandwidth of 4.0 nm, range of 300–1,350 nm, scan speed of 1,000 nm/min. The optical density at (6,5) E_11_ was adjusted to 0.15 to avoid inner filter effects in the fluorescence measurements. Fluorescence emission spectra of nanotubes were acquired by NS Super (Applied NanoFluorescence, Houston TX, USA). The samples were excited at 575 nm (bandwidth of 10 nm, 10-20 mW, averaging: 3, integration time: 2 seconds). Following the acquisition, the data were processed using custom MATLAB codes that applied the spectral corrections and background subtraction, and the fluorescence emission peaks were fitted with Lorentzian functions.

### Confocal Raman Microscopy

Solid state nanotube samples were prepared from aqueous dispersions prior to Raman characterization. Briefly, 100 µL of nanotube suspension (optical density of 1 at (6,5) E_11_) dispersed in sodium deoxycholate (1 wt%, aq) was combined with 1900 µL of ethanol to destabilize the surfactant shell and promote nanotube aggregation. The mixture was centrifuged at 13,000 x g for 5 minutes, after which the supernatant was discarded. The remaining pellet was deposited onto clean glass substrates and dried overnight in a fume hood at room temperature to yield a solid-state film for Raman measurements. Raman spectra were acquired on a QONTOR Confocal Raman spectrometer (Renishaw) under 785 nm excitation laser. Spectra were collected using 1200 lines mm^-1^ diffraction grating, an incident laser power of 40 mW, a 1s integration time, and 25 accumulations per spectrum, with the excitation laser beam focused onto the sample through a 50x objective. Spectra were recorded over the 830–1,900 cm^-1^ spectral acquisition range, measuring both the D band (∼1,300 cm^-1^) and the G band (∼1,590 cm^-1^) of the carbon nanotube samples. Multiple spatial positions across each sample were measured to account for any film heterogeneity. Background subtraction was performed in WiRE 5.2 software (Renishaw) using the standard baseline correction modality. The D/G intensity ratio was calculated from the baseline corrected spectra to quantify relative defect densities of the nanotube samples.

### Zeta Potential Measurements

Aqueous solution of polymer-QWNTs in nanopure water with an optical density at (6,5) E_11_ of 0.15 was transferred into a DTS1070 folded capillary Zetasizer Cell (Malvern, M00049654). Zeta potential values were acquired with the Malvern Zetasizer Nano Z. Instrument parameters for our dispersant are as follows: 25 °C, 0.8872 cP viscosity, refractive index of 1.330, and 78.5 dielectric constant. Equilibrium time is set to 120 seconds. The zeta potentials were calculated with the Smoluchowski model.

### Protein titration experiments

Lyophilized proteins were reconstituted in 1X PBS. The protein sample was transferred to an Amicon Ultracel 3K centrifugal filter (Lot: 0000404176) and centrifuged at 10,000 g until the volume was reduced by 10-fold. The addition of 1X PBS and centrifugation was repeated 5 times. The molarity of the solution was known based on the total amount of protein in the sample and the final volume after dilution in 1X PBS. A stock solution of 2 µM was prepared and subsequent serial dilutions were done to make solutions of 200 nM, 2 nM, 200 pM, and 2 pM. The pooled human plasma was thawed and centrifuged in the same manner. In Eppendorf Protein LoBind Tubes, triplicate samples were prepared in which 14 µL of the plasma supernatant was combined with 7 µL 10x PBS, 35 µL of the appropriate protein stock, nanotube stock, and water to form three 70 µL samples of 0.15 optical density at (6,5) E_11_. Samples were prepared for each of the protein spike concentrations. After 120 min incubation on a shaker, fluorescence measurement is taken as described above.

### Dose-response classification of protein titration experiments

Fluorescence spectra were acquired at each concentration, and changes in four spectral features, E_11_ and E_11_^-^ peak wavelength and intensity, were quantified relative to the 0 µM control. Dose–response curves were modeled as sigmoidal and assigned to near-baseline, linear, or silent by non-parametric criteria. A series was classified as linear if Spearman’s rank correlation between concentration and response was significant (p <0.05). The sign of the Spearman coefficient, ρ, indicates the direction of modulation. A series that did not meet the linear criterion was classified as near-baseline if the triplicate responses at the highest concentration (1 µM) differed significantly from the mean of the responses at all lower concentrations based on a Welch’s t-test. A significant difference (p < 0.05) indicated that the curve captured the onset of the baseline-to-linear transition, where the protein begins to modulate fluorescence only at the highest tested concentration. Spectral responses also required a minimum change of >0.1 nm and >3% intensity change relative to baseline. The remaining series were classified as silent and excluded from subsequent analysis.

### Estimation of half-maximal response concentrations

For series with a significant concentration-dependent response (Dose-response classification, above), the response was summarized by fitting a three-parameter Hill model with the Hill coefficient fixed at unity, in GraphPad Prism (v11.0.0; “[Agonist] vs. response - three parameters”).

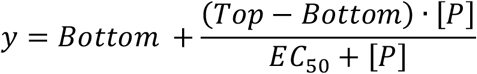

Bottom and Top were left unconstrained, the amplitude (Top - Bottom) was unconstrained in sign so that a single model described both increasing and decreasing responses, and EC_50_ was constrained to positive values. Because majority series, including all near-baseline series, reached a response plateau within the tested range, the upper plateau and the inflection were frequently weakly constrained. The fitted EC_50_ is therefore reported as an apparent half-maximal response concentration rather than as a binding constant. An EC_50_ was reported as a point estimate when its fitted value fell within the tested concentration range (logEC_50_ < −5.85, i.e., EC_50_ < 1.4 µM); series with a fitted EC_50_ above the tested range were reported as lower bounds, and series without a significant response were not fit.

### Protein quantification

The Detergent Compatible Bradford Assay Reagent (Thermo Scientific, Lot: ZE390357) was warmed to room temperature before use. An albumin standard of 2.0 mg/mL in 0.9% NaCl (Thermo Scientific, Lot: ZE392797) was used to prepare a range of standard concentration samples via serial dilution in ultra-pure water as instructed. The standard curve and the protein samples are plated in volumes of 150 µL in a flat-bottom Corning 96 well plate. 150 µL of the Bradford reagent was dispensed rapidly into each well. After 5 minutes of incubation at room temperature and in the dark, the absorbance at 595 nm was read in a Biotek Synergy HT Multi-Mode Microplate Reader.

### MANOVA variance partitioning and permutation testing

To quantify the relative contributions of polymer identity, defect chemistry, and their interaction to both corona composition and fluorescence response, we applied MANOVA modeling with categorical predictors for polymer type and defect type, including a full polymer-QWD interaction term. Variance was partitioned using sequential sum of squares: the polymer main effect was fit first, followed by the defect main effect, then the interaction. The fraction of total variance attributed to each term was computed as the ratio of the explained sum of squares for that term to the total sum of squares. Statistical significance was assessed by permutation testing (5,000 permutations): polymer and defect labels were independently permuted to generate null distributions for main effects, and a Freedman-Lane scheme was used for the interaction term. For proteomics, the model was applied to the 289-protein log2 fold-change matrix using triplicate measurements per construct. To quantify the relative contributions of polymer and QWD to the fluorescence variance, the model was applied separately to wavelength and intensity responses for each emission band, as combining these metrics into a single multivariate response produces scale-dependent weighting artifacts due to their different physical units.

### Tryptic Digestion

All samples were dried via SpeedVac before using the Micro S-Trap (Protifi) columns for tryptic digestion as per the manufacturer’s instructions. Samples were resuspended in 23 µL of 1x sodium dodecyl sulfate Lysis buffer (5% sodium dodecyl sulfate, 50 mM TEAB) and shaken at 2000 rpm for 10 min to lyse the samples and dissolve proteins. Samples were reduced with TCEP to a final concentration of 5 mM and incubated at 55°C for 15 minutes. Reduced samples were then alkylated by the addition of Iodoacetamide (IAA) to a final concentration of 10 mM in solution and incubated at room temperature in the dark. Samples were acidified by the addition of 27.5% phosphoric acid to a final concentration of approximately 2.5% and vortexed to mix. 165 µL of binding buffer (100 mM TEAB (final) in 90% methanol) was added to each tube and shaken at 2000 rpm for 5 mins to mix. The contents of each tube were transferred to an individual S-Trap column placed in a 1.7 mL receiver tube. These tubes were spun at 4000 × g for 1 minute and the flow through was discarded. Trapped proteins were washed three times with wash buffer (100 mM TEAB (final) in 90% methanol), with the flow-through of each wash step discarded. MS grade trypsin protease (Thermo Scientific) was added to each trap column in a 1:50 trypsin to protein ratio dissolved in 20 µL of 50 mM TEAB. The S-Trap columns were incubated in a water bath at 37°C overnight. On the following day, digested peptides were eluted sequentially by 50 mM TEAB, 0.1% formic acid, and 50% acetonitrile. Pooled eluted peptides were dried via SpeedVac. Samples were resuspended in 25 µl of 5% acetonitrile, 0.1% formic acid, shaken at 2000 rpm for 10 minutes, centrifuged at 21 kg for 10 minutes, and the supernatant was collected for LC-MS data acquisition.

### Tandem Mass Tag (TMT) Label Tagging

A TMT 18plex 0.5mg Mass Labeling Kit (Thermo Scientific) was used to isobarically label peptide samples per the manufacturer’s instructions. Immediately before using, the TMT label reagents were equilibrated to room temperature. 20 µL of anhydrous ACN was added to each tube to dissolve the reagent with occasional vortexing for 5 minutes. The protein digest was then resuspended in 10 µL of 100 mM TEAB. The contents of the TMT Reagent vial were transferred to the tubes containing the protein digest in solution. Samples were incubated at room temperature for 1 hour. To quench the reaction, 5 µL of 5% hydroxylamine was added to each sample and incubated at room temperature for 15 minutes. All labeled samples were combined, vortexed and divided into three microcentrifuge tubes for drying via SpeedVac. A pooled plasma reference channel was included to enable quantitative fold-change analysis.

### Reversed Phase-Reversed Phase Fractionation

A High pH Reversed-Phase Peptide Fractionation Kit (Thermo Scientific) was used to fractionate TMT-labeled peptides per the manufacturer’s instructions. The spin columns were placed into a collection tube and conditioned by centrifugation at 5,000 g for 2 minutes to remove the storage solution and pack the resin material. To wash the columns, 300 µL of anhydrous acetonitrile was added, and the spin column was centrifuged at 5,000 g for 2 minutes. This wash step was repeated with the flow-through of the wash discarded. The column was then washed twice, as in the previous step, with 0.1% TFA to complete the column conditioning. Elution solutions were prepared according to Supplementary Table 1. Labeled proteolytic digests were dissolved in 100 µL of 0.1% TFA and combined to a total of 300 µL. The conditioned spin column was placed into a new 2.0 mL receiver tube, and the 300 µL resuspended protein digest was loaded onto the column. The column was centrifuged at 3,000 g for 2 minutes, and the flow-through was retained. This step was repeated with water loaded onto the column, and the flow-through was collected as the “first wash” fraction. The “second wash” fraction was collected using the first elution solution of Supplementary Table 6 to remove unreacted TMT reagent. The column was placed in a new receiving tube and 300 µL of the remaining gradient elution solutions from Supplementary Table 6 were used with each fraction being collected by centrifugation at 3,000 g for 2 minutes. Each fraction was dried via SpeedVac and the dried samples were resuspended in 25 µL of 0.1% formic acid, shaken at 2,000 rpm for 10 minutes, and centrifuged at 21 kg for 10 minutes before being placed in sample vials of 15 µL for LC-MS analysis.

### LC-MS/MS Data Acquisition

An externally calibrated Thermo Exploris 480 (high-resolution electrospray tandem mass spectrometer) was used in conjunction with Vanquish Neo nano LC System. 1 μg of each sample was aspirated into a 50 μL loop and loaded onto the trap column (Thermo µ-Precolumn 5 mm, with nanoViper tubing 30 µM i.d. × 10 cm). The flow rate was set to 300 nL/min for separation on the analytical column (EASY-Spray™ PepMap™ Neo UHPLC Column, 50 cm long, 75-micron internal diameter, C18, reverse phase). Mobile phase A was composed of 99.9% H2O (EMD Omni Solvent), and 0.1% formic acid and mobile phase B was composed of 80% ACN, and 0.1% formic acid. A 90-minute linear gradient from 5% to 55% B was performed. The LC eluent was directly nanosprayed into Exploris 480 mass spectrometer (Thermo Scientific). During the chromatographic separation, the Exploris 480 plus was operated in a data-dependent mode and under direct control of the Thermo Excalibur 4.7.69.37 Software (Thermo Scientific). MS data was acquired using the following parameters: data-dependent scan with a 3 second cycle time per full scan (400 to 1500 m/z) at 60,000 resolutions. MS2 was acquired at 15,000 resolutions. Ions with a single charge or charges of more than 8, as well as unassigned charges, were excluded. An auto-dynamic exclusion window was used. All measurements were performed at room temperature.

### LC-MS/MS Data Analysis

Resultant raw files were searched with Proteome Discoverer 3.2 using the Chimerys search engine with a *Homo sapien* FASTA database. A 20-ppm mass tolerance for parent ion and 0.02Da mass tolerance for the fragment ion was used.

