## Supplementary Information for "Structure-affinity correlations and separable optical activity in carbon nanotube protein coronas"

Supplementary References………………………………………………………………….24

**Supplementary Methods: Reanalysis of Goerzen et al. fluorescence titration data**

We reanalyzed the fluorescence titration data reported by Goerzen et al.[1] for DSPE-PEG wrapped 3,4,5-trifluoroaryl QWNTs (Supplementary Figs 16-18). For the enrichment comparison, the 22 proteins present in both the Goerzen dataset and our corona proteomics were grouped as enriched (n = 11) or depleted (n = 11) by their corona enrichment (Supplementary Figs 16-17). The dataset comprised triplicate measurements for plasma proteins across seven concentrations (0.015–246 µg/mL) and four spectral features per protein: the $E_{11}$ and $E_{11}^{-}$ wavelength shifts (*dwl*, *dwl** in nm) and the $E_{11}$ and $E_{11}^{-}$ intensity changes (*dint*, *dint** in %). Individual replicate values were used throughout the analysis.

Molecular weights for the protein standards used in the titrations were taken from the source and vendor information reported in the original study, and each protein was converted from mass concentration to molar concentration. For every protein and spectral feature, the response was offset relative to the mean of its triplicate at the lowest tested concentration (0.015 µg/mL), so that the lowest concentration defined the zero baseline. All subsequent steps used this relative response.

To match the accessible concentration range of our own titration, which extended to 1 µM, the analysis window for each protein was capped at the concentration nearest 1 µM in log space; that concentration and all lower concentrations were retained, and higher concentrations were excluded. Because molecular weights differ among proteins, five to seven concentrations were retained for each protein.

Each protein-feature response was classified on the retained window. A response was called linear if Spearman's rank correlation between concentration and relative response was significant ($p < 0.05$); the sign of the correlation gave the direction. A response that was not linear was called near-baseline if the relative responses at the highest retained concentration differed from those at the lower retained concentrations by Welch's t-test ($p < 0.05$) and a concordant monotonic trend was confirmed by the Jonckheere-Terpstra test in the direction of the top-concentration change. This monotonicity requirement excluded single-concentration deviations that the t-test alone would admit. A called response was additionally required to exceed a magnitude floor at the highest retained concentration, set at 0.1 nm for the wavelength features and 5% for the intensity features. Responses below the cutoffs were treated as not significant. Responses meeting neither the linear nor the near-baseline criterion were classified as silent. For each response, a three-parameter Hill model was fit to the retained points by nonlinear least squares as described in our Methods section.

**Supplementary Note 1: Descriptor-resolved effects of QWD chemistry on corona composition**

Five chemical descriptors of the aryl QWD were considered with the following physicochemical categories: electronic (Hammett constant), steric (hydrodynamic size), and thermodynamic (H-bond capacity, logP). To distinguish genuine descriptor-specific effects from collinearity artifacts, we focus below on features exclusively significant for one descriptor class.

The electronic character of QWDs correlates with the depletion of positively charged surface residues. More H-donating and hydrophobic QWDs negatively correlate with GRAVY and positively with exposed aromatics, indicating a preference for hydrophilic proteins with high surface exposed aromatics. At the amino acid level, electron-withdrawing QWDs uniquely enrich methionine and deplete lysine, indicating a preference for uncharged protein surfaces. H-bond accepting QWDs enrich strong H-bond donors, e.g., glutamine, while H-bond donor QWDs preferentially recruit acceptor-capable surface residues. Surface serine, phenylalanine, and tyrosine residues, which can accept hydrogen bonds or participate in hydrogen-pi interactions, positively correlate with this axis.

Structural features, including flexibility and secondary structure information, are significant for both Hammett and the thermodynamic axis, with opposite correlations. However, because Hammett and the thermodynamic axis are anti-correlated, these structural features cannot be attributed to either descriptor independently. Resolving this conformational selection trend would require an expanded library in future work.

In terms of steric space, bulkier QWDs recruit larger, more polar corona proteins, but does not significantly correlate with any structural or flexibility metric. This differentiates the steric axis related to protein size and polarity selection from the conformational effects captured by the electronic/thermodynamic gradients described in the main text.

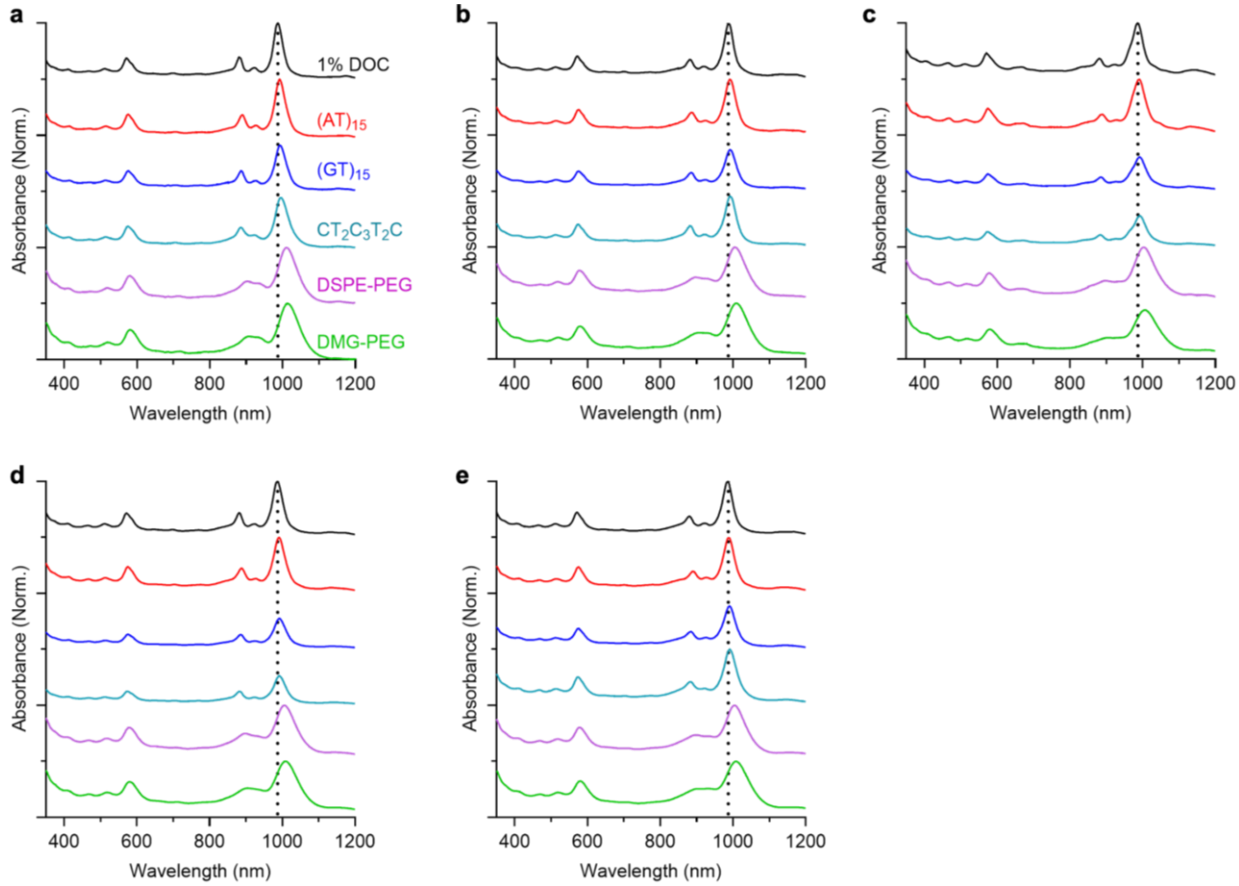

**Supplementary Figure 1.** Absorbance spectra of chirality separated QWNT stocks before polymer exchange and after rewrapping with $(AT)_{15}$, $(GT)_{15}$, $CT_2C_3T_2C$, DSPE-PEG, DMG-PEG. Spectra are normalized to (6,5) $E_{11}$ absorbance band. Dotted vertical line centered on (6,5) $E_{11}$ center wavelength in 1% DOC (aq.) to visualize red shifting of polymer wrapped nanotubes. a, Pristine, b, Amino, c, Dinitro, d, Carboxyl, e, Trifluoro.

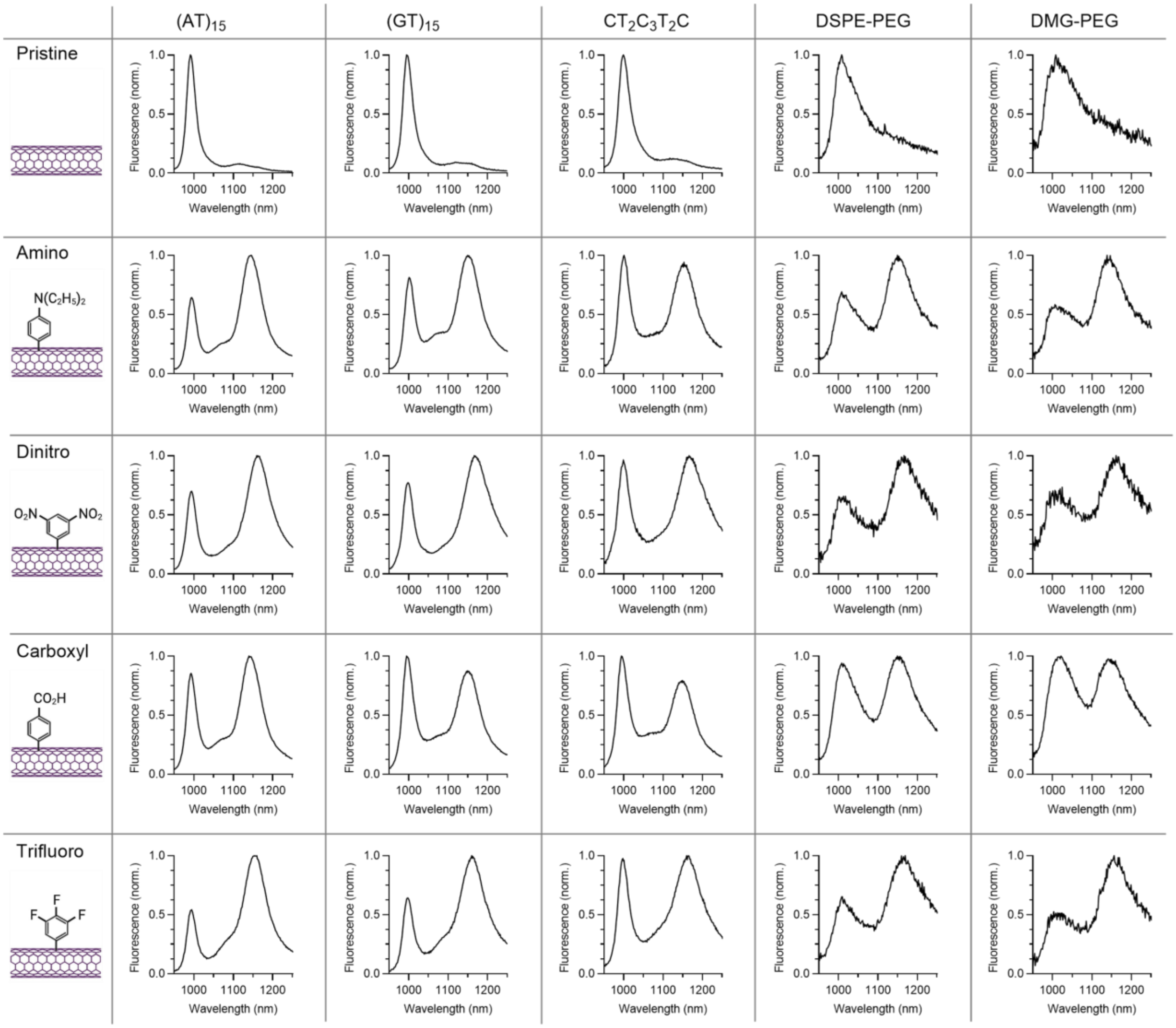

**Supplementary Figure 2.** Normalized fluorescence spectra of nanotube stock solutions in 1x PBS. The concentration of (6,5)-SWCNTs is 2 ug/mL. Rows indicate QWD identity; columns indicate polymer identity.

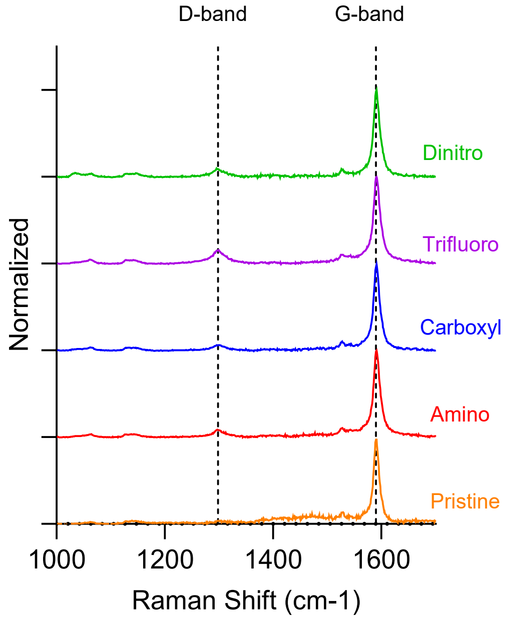

**Supplementary Figure 3.** Raman spectra of pristine and QWD-functionalized nanotubes. Full Dashed lines indicate Raman D and G bands. Each spectrum was acquired with 40 mW of 785 nm laser excitation and normalized with respect to the G-band.

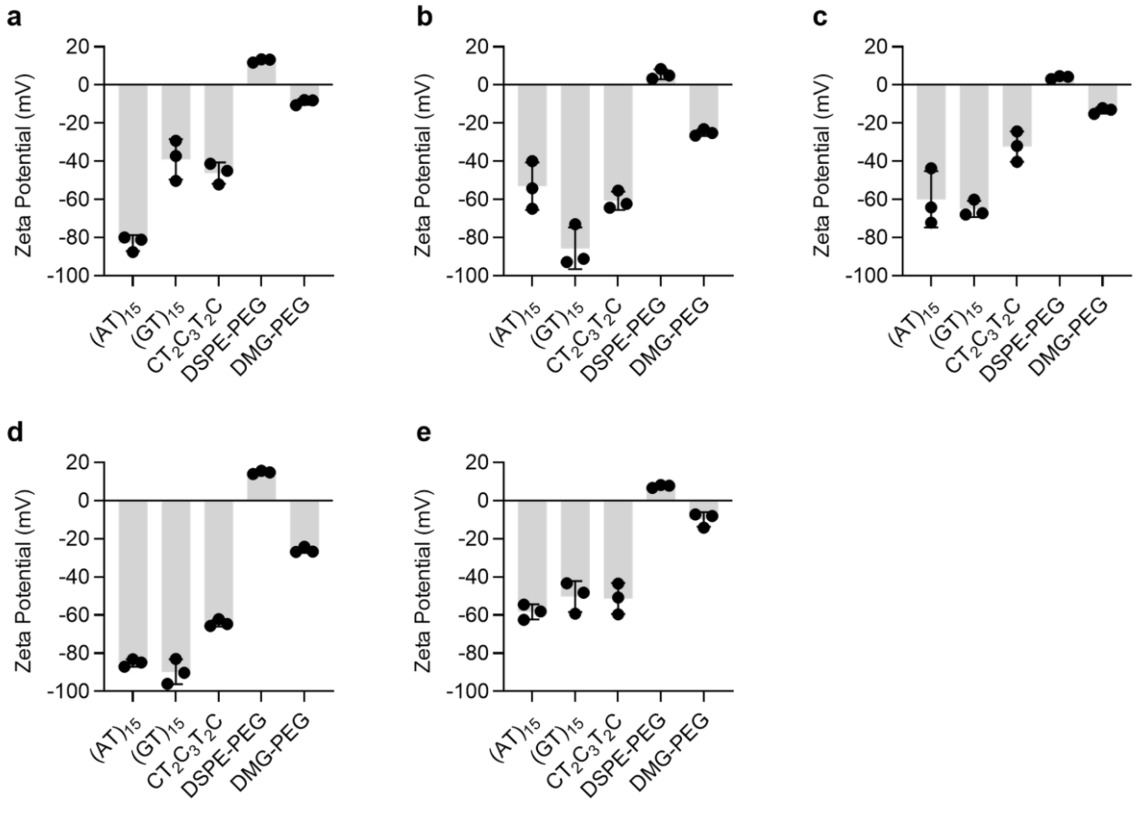

**Supplementary Figure 4.** Zeta potential of polymer-wrapped QWNTs at 0.15 OD of (6,5) $E_{11}$ in DI water. Error bars represent **the** standard deviation of triplicates. **a,** Pristine, **b,** Dinitro, **c,** Amino, **d,** Carboxyl, **e,** Trifluoro.

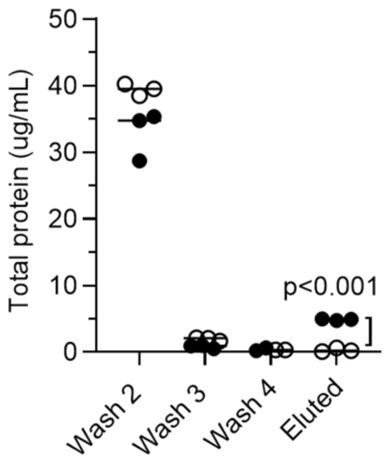

**Supplementary Figure 5.** Bradford quantification of protein in the 1x PBS wash supernatants and in the eluate from the nanotube pellet. Filled circles are the SWCNT included triplicates; empty circles are SWCNT-excluded controls.

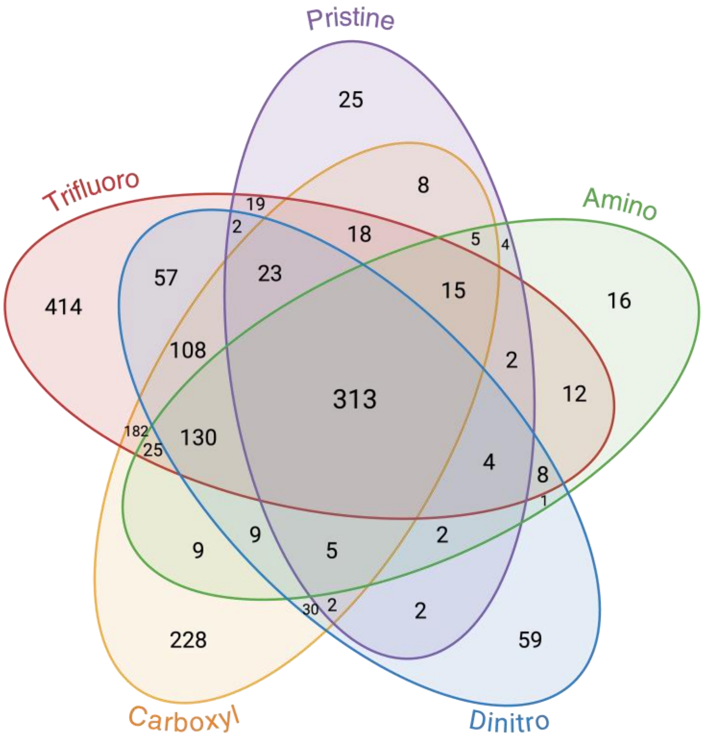

**Supplementary Figure 6.** Five-way Venn diagram of proteins identified across the nanotube coronas. Values indicate the number of proteins shared among the indicated coronas.

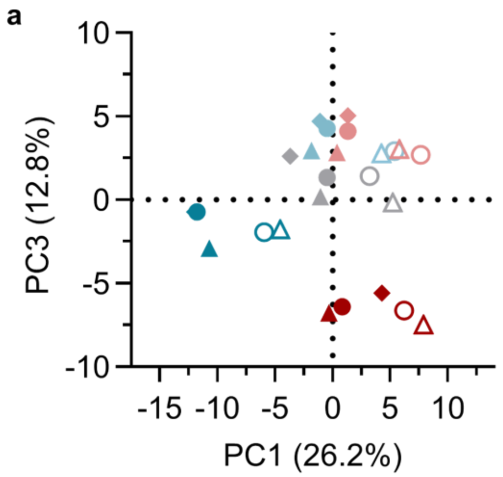

**Supplementary Figure 7.** PCA plot of all sensor coronas, derived from Log2 protein fold-changes in 289 common proteins. Red, light red, gray, light blue, and blue symbols indicate dinitro, carboxyl, trifluoro, and aminoaryl QWNTs and pristine nanotubes, respectively. The filled triangle, circle, and diamond are $CT_2C_3T_2C$, $(GT)_{15}$, and $(AT)_{15}$ wrapping, respectively. The empty triangle and circle are DMG-PEG and DSPE-PEG wrapping, respectively. **a,** PC1 versus PC3. **b,** PC2 versus PC3.

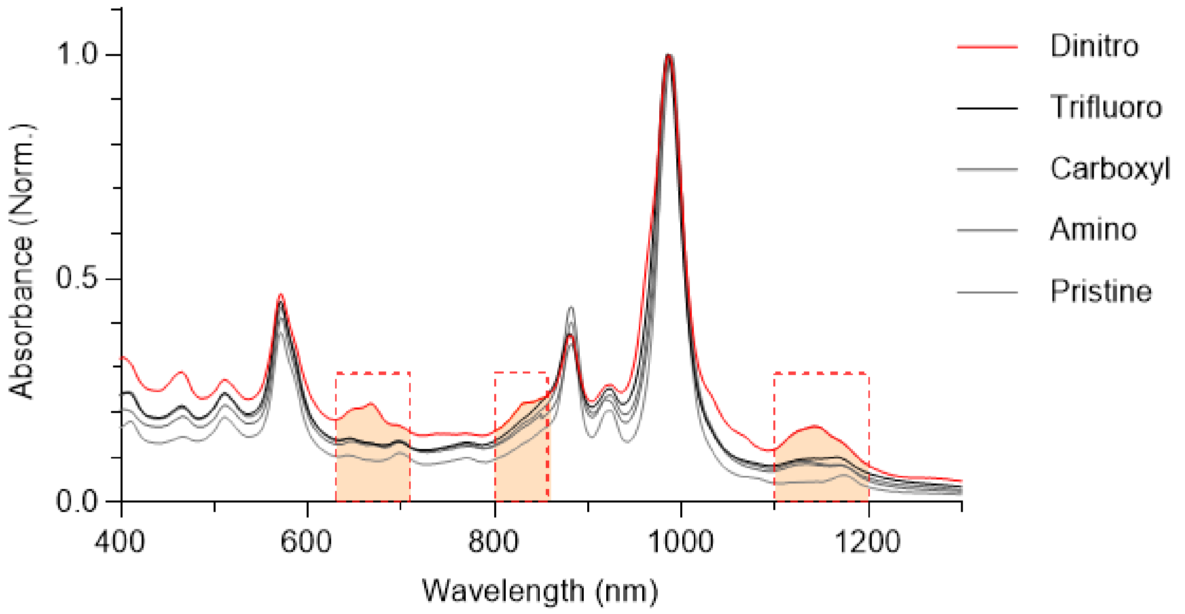

**Supplementary Figure 8.** Normalized, overlaid absorbance spectra of nanotube stock in 1% DOC (aq.). Highlighted regions in dashed red boxes mark $E_{11}$ and $E_{22}$ absorption bands from non-(6,5) chiralities observed in 3,5-dinitroaryl QWNTs.

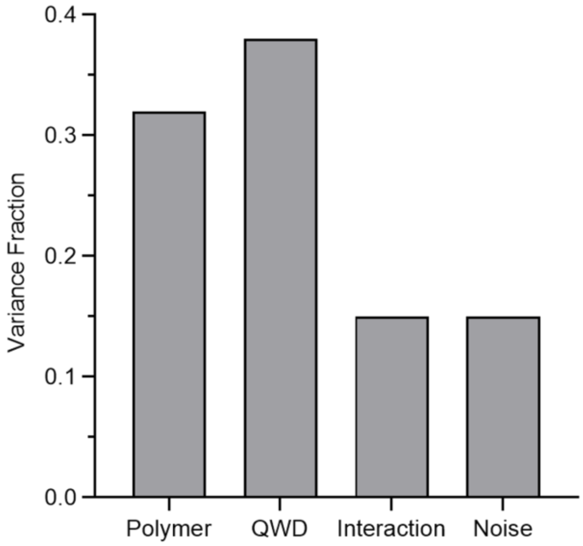

**Supplementary Figure 9.** Partitioned variance in corona compositions for common proteins, excluding pristine coronas.

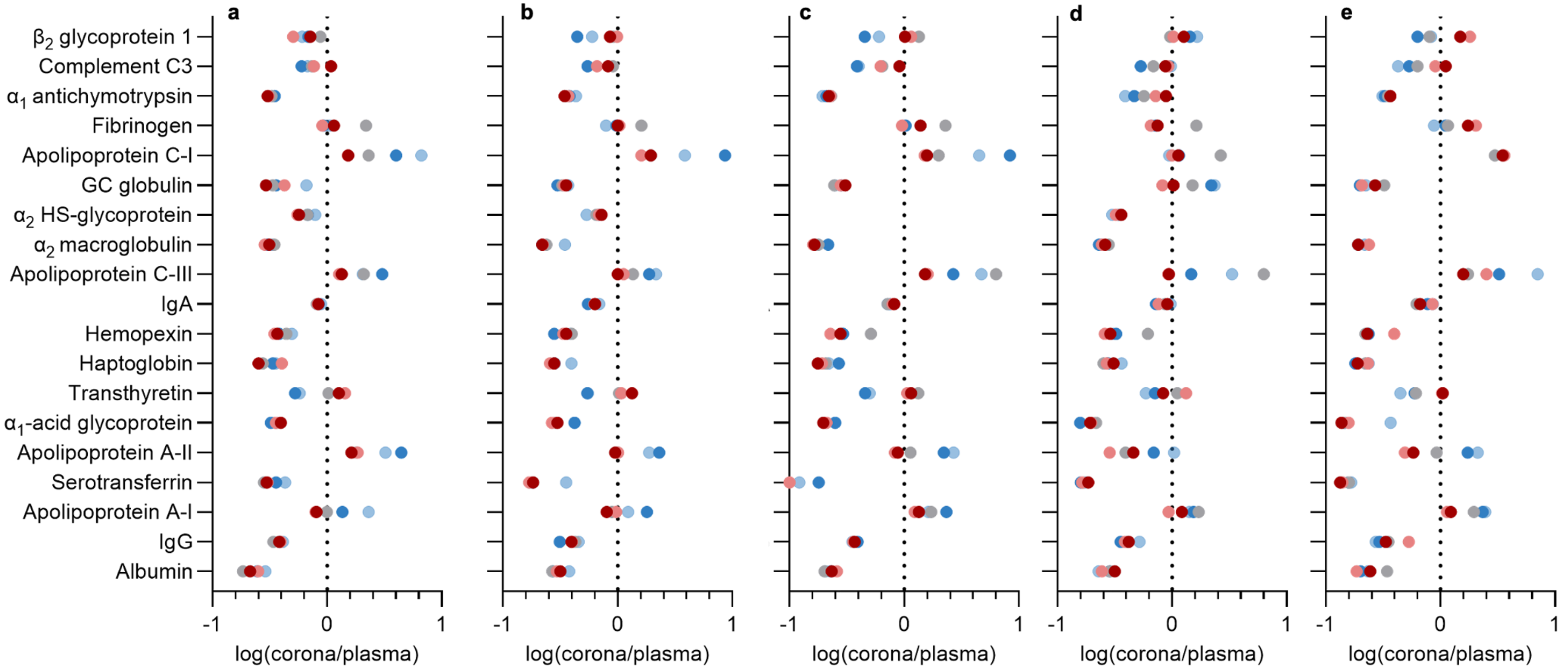

**Supplementary Figure 10.** $Log_2$ fold-change plots of the 19 most abundant plasma proteins by molarity in each corona relative to plasma. **a,** Pristine, **b,** Amino, **c,** Dinitro, **d,** Trifluoro, **e,** Carboxyl. Blue: DSPE-PEG, Light blue: DMG-PEG, Gray: $CT_2C_3T_2C$, Light red: $(AT)_{15}$, Red: $(GT)_{15}$.

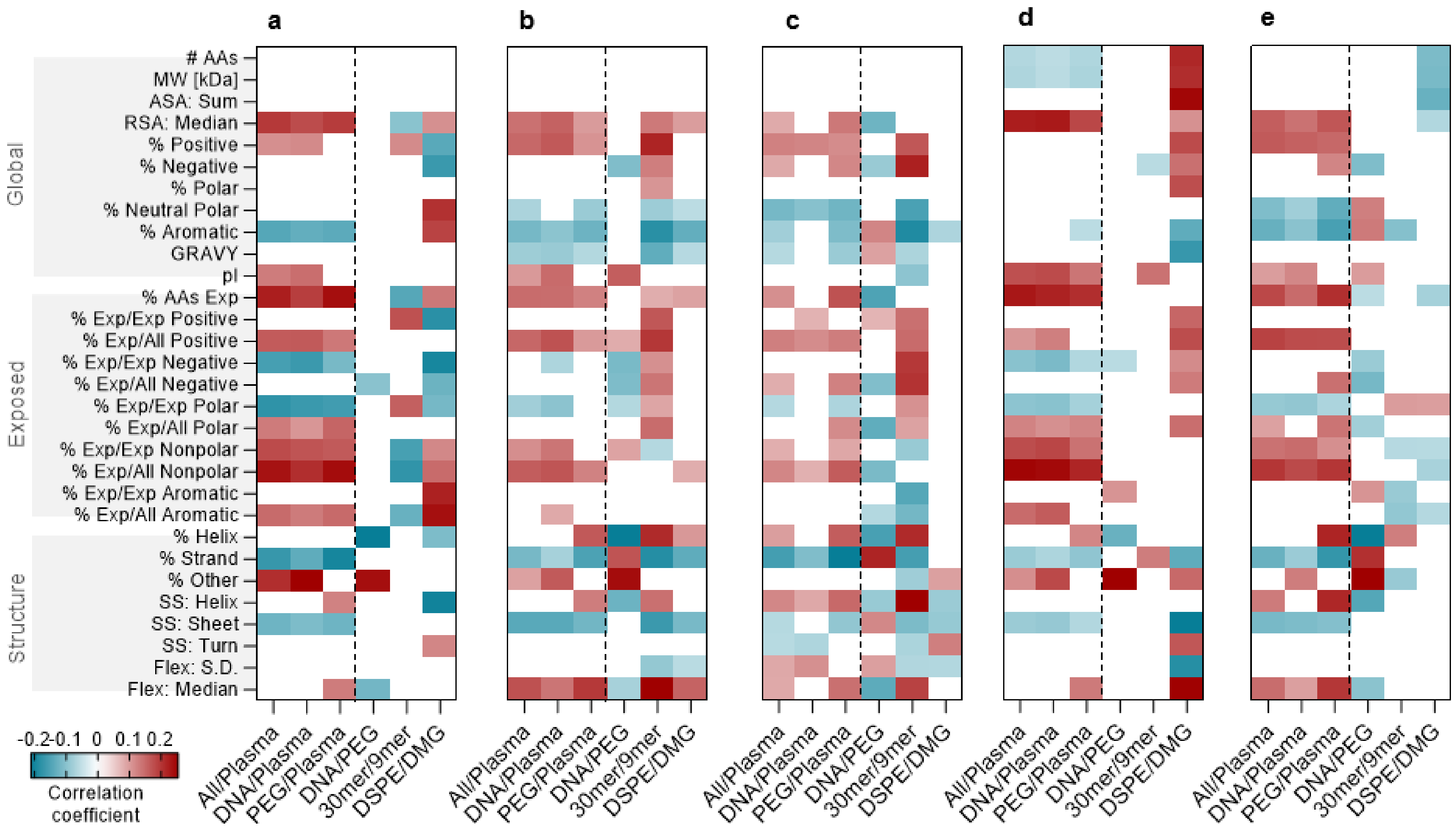

**Supplementary Figure 11.** Heatmaps of Pearson rho values for correlations between protein features and corona fold change. **a,** Pristine, **b,** Amino, **c,** Dinitro, **d,** Trifluoro, **e,** Carboxyl.

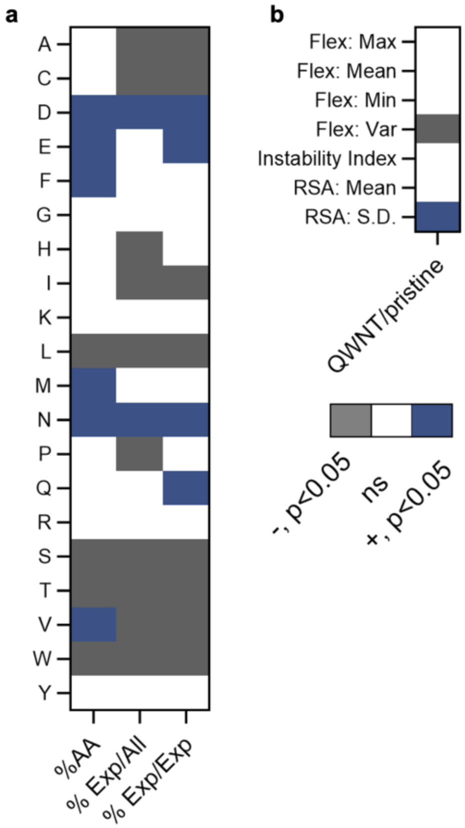

**Supplementary Figure 12.** Heatmaps of Mann-Whitney test values comparing weighted protein features between QWNT and pristine nanotubes. **a,** Per-residue amino acid content, including %AA, % Exp/All, and % Exp/Exp. **b,** Flexibility, instability, and surface area features. Color indicates the direction of the significant ($p<0.05$) difference. Gray: Lower in QWNT; white: Not significant; Blue: Higher in QWNT.

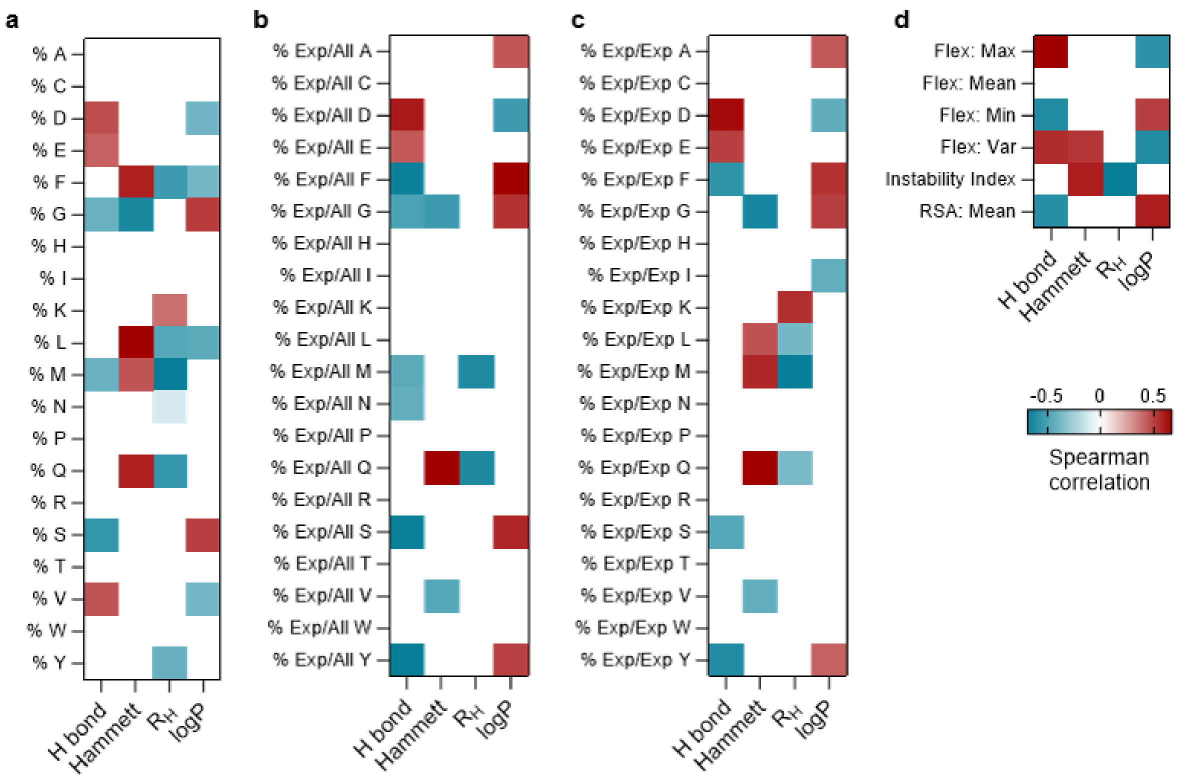

**Supplementary Figure 13.** Heatmaps of Spearman rho values between weighted protein features and chemical parameters of QWD. **a,** %AA. **b,** % Exp/All. **c,** % Exp/Exp. **d,** Flexibility, instability, and surface area features. RSA: Relative surface area. Only significant correlations ($p<0.05$) are shown.

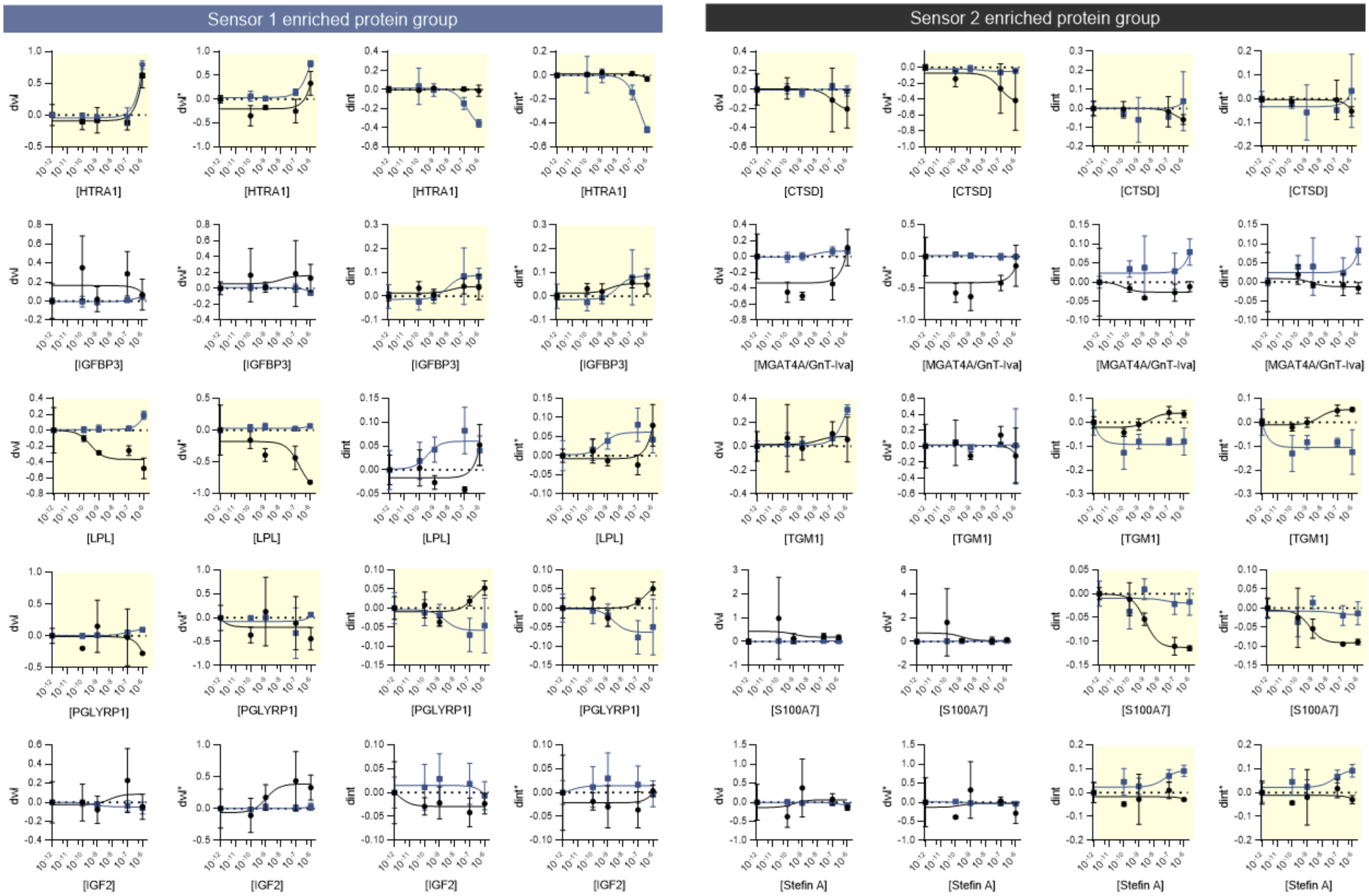

**Supplementary Figure 14.** Sensor response in 20% plasma. Blue: Sensor 1 with Trifluoro QWD and $(AT)_{15}$ wrapping; Black: Sensor 2 with Amino QWD and DSPE-PEG wrapping. Yellow background marks a statistically significant spectral response. Curves are three-parameter Hill fits and guide the eye. Error bars give the standard deviation of technical replicates.

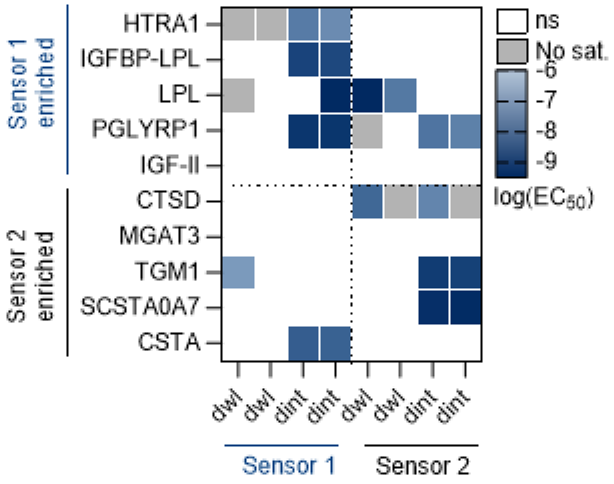

**Supplementary Figure 15.** Heatmap of log-transformed $EC_{50}$ values for sensor response across ten proteins and four spectral features per sensor. For active responses, apparent half-maximal response concentrations ($EC_{50}$) were estimated from three-parameter Hill fits. Color indicates log-transformed $EC_{50}$ values in analyte concentration. Gray cells indicate the spectral responses did not saturate within the tested titration range and $EC_{50}$ > -6.

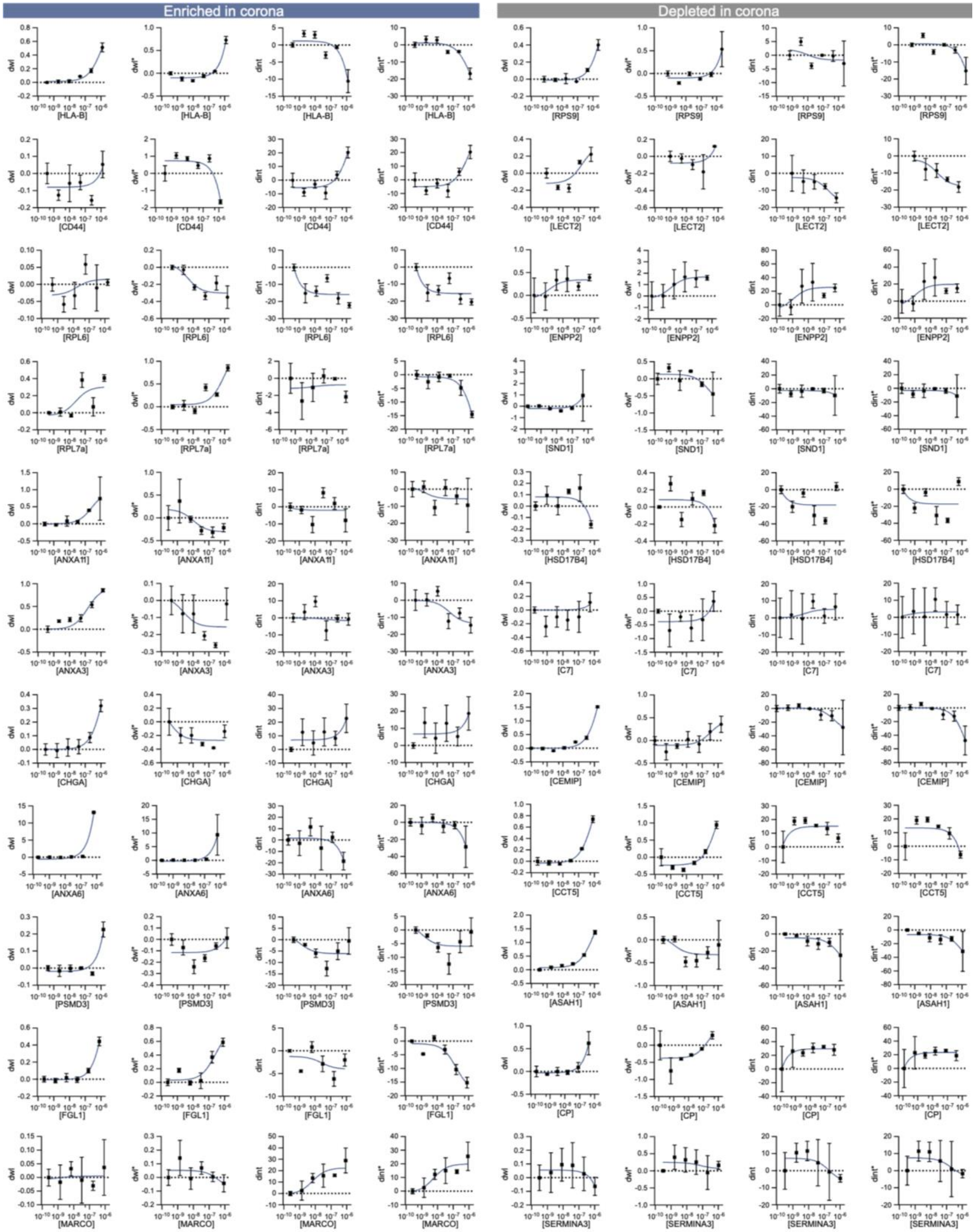

**Supplementary Figure 16.** Fluorescence response of DSPE-PEG wrapped 3,4,5-trifluoroaryl QWNTs in 20% plasma. Fluorescence data were from the previous report by Goerzen et al.[1] Curves are three-parameter Hill fits and guide the eye. Error bars: Standard deviation.

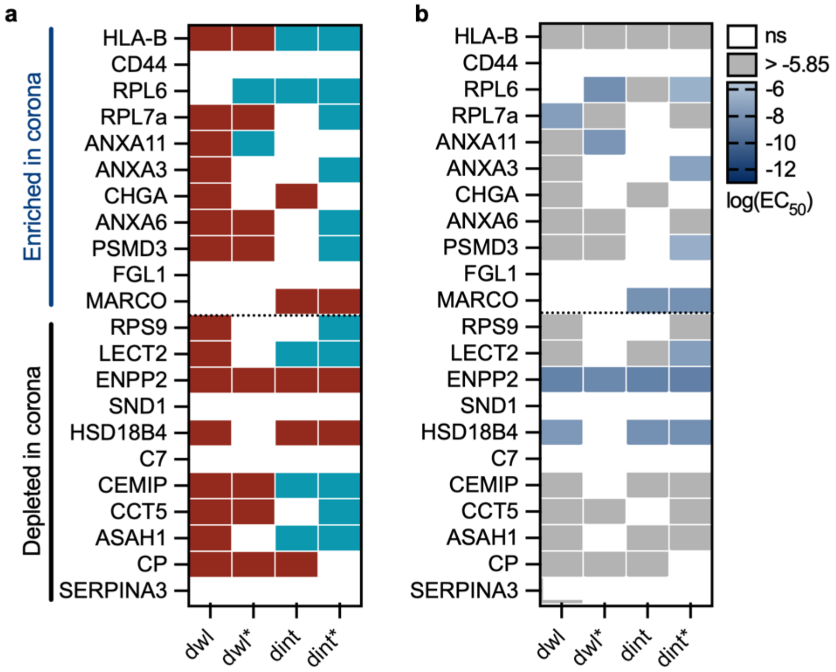

**Supplementary Figure 17.** Heatmap of sensor response across an independent protein panel. Reanalysis of the fluorescence titration data of Goerzen et al. for DSPE-PEG wrapped 3,4,5-trifluoroaryl QWNTs, for the 22 proteins shared with our corona proteomics. Proteins are grouped as enriched (top) or depleted (bottom) in the corona. Responses were called from triplicate titrations over the ≤1 μM window. **a,** Direction of the significant responses. Red and blue cells indicate an increase and decrease of significant spectral response. White: not significant. The nanotube responded to enriched and depleted proteins at comparable rates (9/11 vs. 8/11). **b,** Half-maximal response concentration from three-parameter Hill fits. Color indicates log-transformed $EC_{50}$ values in analyte concentration. Gray cells indicate the spectral responses did not saturate within the tested titration range and $EC_{50} > 6$.

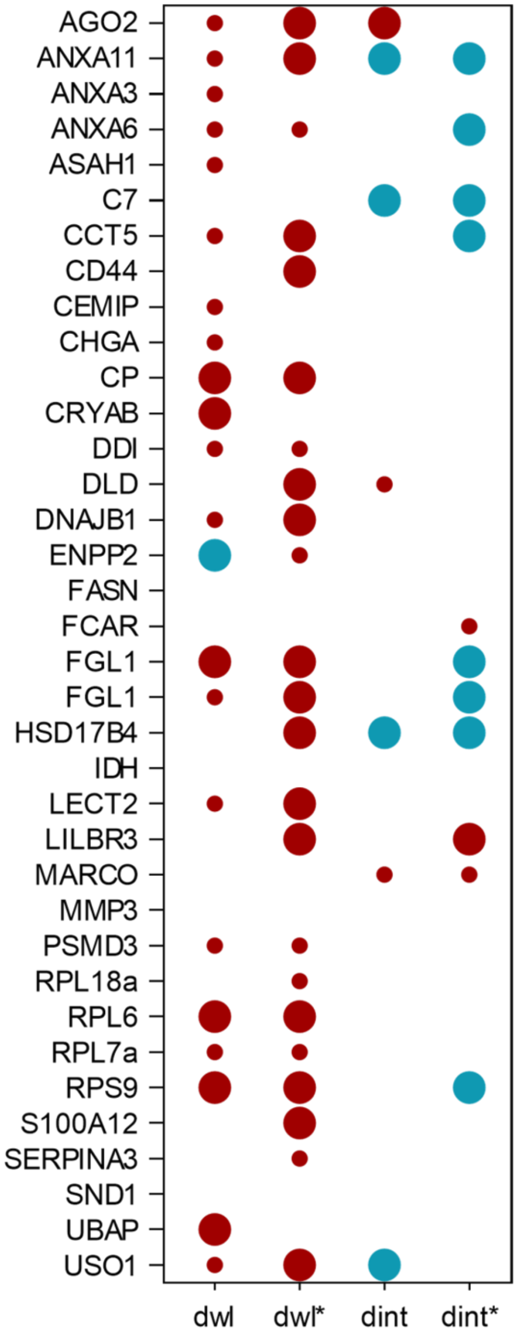

**Supplementary Figure 18.** Fluorescence response of all DSPE-PEG wrapped 3,4,5-trifluoroaryl QWNTs proteins. Fluorescence titration data from Goerzen et al.[1] for the full protein panel, listed alphabetically. Each protein was titrated from 0.015 to 246 μg/mL. Circles mark significant responses; color gives direction (red, increase; blue, decrease) and size gives regime (small, near-baseline; large, linear). Only responses significant above a magnitude cutoff of 1 nm for the wavelength features and 20% for the intensity features are presented.

**Supplementary Table 1.** Chemical parameters of QWDs in buffer protonation states. Six chemical descriptors of the aryl QWD categorized with the following physicochemical variable: electronic (Hammett constant), steric (polarizability, relative volume, hydrodynamic size), and thermodynamic (H-bond capacity, logP). Within the steric category, polarizability, relative volume, and hydrodynamic size ($R_H$) are nearly interchangeable, so we used $R_H$ as a single representative steric descriptor in the main text analysis. H-bond capacity and logP are rank-correlated across the QWD series, and we group them as a single thermodynamic axis.

Relative volume: McGowan volume ($cm^3$/mol). $R_H$: Hydrodynamic radius derived using the standard Wilke-Chang Equation for diffusion in water, then converted diffusivity into hydrodynamic radius using the Stokes-Einstein equation. Hammett: Hammett substituent coefficient of terminating moieties of aryl QWDs. H-bond: Categorical rank of hydrogen bond donor capability of QWDs at pH 7.4 (high to low: H-bond acceptor to donor). Polarizability: Molar refractivity ($cm^3$/mol).

| Terminating group of aryl QWD | Relative volume | $R_H$ | Polarizability | Hammett | H-bond | logP | Charge |
|---|---|---|---|---|---|---|---|
| N,N-diethyl-4-amino | 137.98 | 3.48 | 49.8 | -0.83 | 1 | 3.7 | 0.126 |
| 3,5-dinitro | 106.48 | 2.83 | 37.6 | 1.42 | 3 | 1.5 | 0 |
| 4-carboxyl | 93.17 | 3.04 | 31 | 0 | 4 | -2 | -0.999 |
| 3,4,5-trifluoro | 76.95 | 2.44 | 26.1 | 0.74 | 2 | 2.6 | 0 |

**Supplementary Table 2.** Protein features analyzed in this study.

| Category | Variable | Description |
|---|---|---|
| Global | # AAs | Number of total amino acids |
| | MW [kDa] | Molecular weight in kDa |
| | ASA: Sum | Absolute surface area |
| | RSA: Median | Relative surface area, median |
| | % Positive | Fraction of positively charged amino acids (R,H,K) |
| | % Negative | Fraction of negatively charged amino acids (D,E) |
| | % Polar | Fraction of polar amino acids (S,T,N,Q,C,Y,D,E,K,R,H) |
| | % Neutral Polar | Fraction of neutral, polar amino acids (S,T,N,Q,C,Y) |
| | % Nonpolar | Fraction of neutral, nonpolar amino acids (G,A,V,L,I,M,F,W,P) |
| | % Aromatic | Fraction of aromatic amino acids (F,Y,W,H) |
| | GRAVY | Grand Average of Hydropathy, a measure of overall hydrophobicity |
| | pI | Isoelectric point |
| Surface | % AAs Exposed | Fraction of exposed amino acids over all amino acids |
| | % Exp/Exp Positive | Fraction of positive exposed amino acids over all exposed amino acids |
| | % Exp/All Positive | Fraction of positive exposed amino acids over all amino acids |
| | % Exp/Exp Negative | Fraction of negative exposed amino acids over all exposed amino acids |
| | % Exp/All Negative | Fraction of negative exposed amino acids over all amino acids |
| | % Exp/Exp Polar | Fraction of polar exposed amino acids over all exposed amino acids |
| | % Exp/All Polar | Fraction of polar exposed amino acids over all amino acids |
| | % Exp/Exp Nonpolar | Fraction of nonpolar exposed amino acids over all exposed amino acids |
| | % Exp/All Nonpolar | Fraction of nonpolar exposed amino acids over all amino acids |
| | % Exp/Exp Aromatic | Fraction of aromatic exposed amino acids over all exposed amino acids |
| | % Exp/All Aromatic | Fraction of aromatic exposed amino acids over all amino acids |
| Structure | % Helix | Fraction of amino acids predicted to be part of an alpha helix |
| | % Strand | Fraction of amino acids predicted to be part of a beta strand |
| | % Other | Fraction of amino acids predicted to be part of a loop instead of an alpha helix or a beta sheet. |
| | SS: Helix | Secondary structure: Alpha Helix |
| | SS: Turn | Secondary structure: Turn |
| | SS: Sheet | Secondary structure: Beta Sheet |
| | Flex: S.D. | Flexibility, standard deviation |
| | Flex: Median | Flexibility, median |

**Supplementary Table 3.** Functionalization via in-situ production of diazonium.

| Aniline | SG65i (mg) | Water (mL) | 10x PBS (uL) | $NOBF_4$ (umol) | Aniline (umol) | $H_2O_2$ (50% wt. in $H_2O$) | Reaction Time |
|---|---|---|---|---|---|---|---|
| 4-aminobenzoic acid<br>Sigma-Aldrich<br>Lot: 1003627281 | | 0.5 | | 4.2 | 4.2 | | |
| | 0.8 | | 100 | | | 200 uL | 3 Days |
| 3,4,5-trifluoroaniline<br>TCI Chemical<br>Lot: 78K8K-IK | | 0.7 | | 4.48 | 4.48 | | |

**Supplementary Table 4.** Diazonium functionalization conditions for each QWNT type.

| Diazonium | SG65i (2mg/mL) | 10x PBS | Diazonium salt | $H_2O_2$ (50% wt. in $H_2O$) | Reaction Time |
|---|---|---|---|---|---|
| N,N-diethyl-4-aminobenzene diazonium tetrafluoroborate | 400 uL | | 0.2 mg | 200 uL | 3 Days |
| | | 100 uL | | | |
| 3,5 dinitrobenzene diazonium tetrafluoroborate | 500 uL<br>+310 uL $H_2O$ | | 0.36 mg | 197 uL | 1 Hour |

Diazonium salts were synthesized following the protocol reported by Piao et al.[2] N,N-Diethyl-p-phenylenediamine (Sigma-Aldrich, Lot: STBK1127) and 3,5-dinitroaniline (Sigma-Aldrich, CAS-No: 618-87-1) were used as received.

**Supplementary Table 5.** Protein specifications. All proteins were purchased from R&D Systems.

| Protein | Catalog no. | Lot no. | Source | Notes |
|---|---|---|---|---|
| Cystatin A | 1407-PI | IGO0224122 | Recombinant human, E. coli derived | 6-His tag, >95% purity |
| IGFBP-3 | 675-B3 | BUY1325061 | Recombinant human, NS0-derived | >95% purity |
| PGLYRP1/PGRP-S | 2590-PGB | NLC1725081 | Recombinant human, NS0-derived | 6-His tag, >95% purity |
| IGF-II/IGF2 | 292-G2 | DS3025051 | Recombinant human, E. coli derived | >97% purity |
| 1/TGM1 | 7765-TG | DCNB0324121 | Recombinant human, Sf 21-derived | 6-His tag, >70% purity |
| HTRA1/PRSS11 | 2916-SE | DAAJ0723121 | Recombinant human, E. coli derived | 6-His tag, >90% purity |
| S100A7 | 9085-SA | QCM0325021 | Recombinant human, E. coli derived | >95% purity |
| Cathepsin D | 1014-AS | FSP0325101 | Recombinant human, NS0-derived | 10-His tag, >95% purity |
| Lipoprotein Lipase | 9888-LL | DISO0525071 | Recombinant human, CHO-derived | 6-His tag |
| N-acetylglucosaminy ltransferase III/MGAT3 | 7359-GT | DCAY0625081 | Recombinant human, CHO-derived | 6-His tag, >90% purity |

**Supplementary Table 6.** Preparation of elution solutions for Thermo-Scientific TMT-labeled peptides.

| Fraction Number | Percent of Acetonitrile |
|---|---|
| Second Wash | 5.0 |
| 1 | 7 |
| 2 | 12 |
| 3 | 17 |
| 4 | 22 |
| 5 | 30 |
| 6 | 50 |
